# Vascular oxidative stress causes neutrophil arrest in brain capillaries, leading to decreased cerebral blood flow and contributing to memory impairment in a mouse model of Alzheimer’s disease

**DOI:** 10.1101/2023.02.15.528710

**Authors:** Nancy E. Ruiz-Uribe, Oliver Bracko, Madisen Swallow, Argen Omurzakov, Sabyasachi Dash, Hiroki Uchida, David Xiang, Mohammad Haft-Javaherian, Kaja Falkenhain, Michael E. Lamont, Muhammad Ali, Brendah N. Njiru, Hsin-Yun Chang, Adrian Y Tan, Jenny Z Xiang, Costantino Iadecola, Laibaik Park, Teresa Sanchez, Nozomi Nishimura, Chris B. Schaffer

## Abstract

**INTRODUCTION:** In this study, we explore the role of oxidative stress produced by NOX2-containing NADPH oxidase as a molecular mechanism causing capillary stalling and cerebral blood flow deficits in the APP/PS1 mouse model of AD.

**METHODS:** We inhibited NOX2 in APP/PS1 mice by administering a 10 mg/kg dose of the peptide inhibitor gp91-ds-tat i.p., for two weeks. We used *in vivo* two-photon imaging to measure capillary stalling, penetrating arteriole flow, and vascular inflammation. We also characterized short-term memory function and gene expression changes in cerebral microvessels.

**RESULTS:** We found that after NOX2 inhibition capillary stalling, as well as parenchymal and vascular inflammation, were significantly reduced. In addition, we found a significant increase in penetrating arteriole flow, followed by an improvement in short-term memory, and downregulation of inflammatory gene expression pathways.

**DISCUSSION:** Oxidative stress is a major mechanism leading to microvascular dysfunction in AD, and represents an important therapeutic target.

## 1. INTRODUCTION

Alzheimer’s disease (AD) is the most common form of dementia, affecting millions of people worldwide [1]. Characterized by the aggregation of amyloid beta plaques in brain tissue and neurofibrillary tangles inside neurons, recent research has highlighted the importance of vascular risk factors [2,3], such as diabetes [4], hypertension [5], and stroke [6], in the development and initiation of the disease. AD pathogenesis is commonly accompanied by vascular abnormalities, such as white matter lesions, cerebral hypoperfusion [7], endothelial and pericyte dysfunction [8], blood-brain barrier leakage [9], and impaired neurovascular coupling [10,11]. Among these vascular abnormalities, cerebral blood flow (CBF) decrease is one of the earliest biomarkers of the disease, and a baseline reduction of ~20% has been detected in AD patients [12] and AD mouse models [5]. In patients with mild cognitive impairment or AD, decreased CBF was found in diverse areas of the brain, including the frontal, temporal, orbitofrontal cortices, and in the hippocampus [7]. In addition, several reports suggest that the severity of CBF reduction in AD patients correlates with the severity of memory impairment and amyloid pathology [13–15]. Although these findings suggest CBF deficits may have a large impact on neurodegeneration and disease progression, the causes of CBF deficits in AD and related dementias constitutes a field of active research [16–20]. Improving our understanding of the molecular and cellular causes of the CBF reduction may provide critical insights into disease mechanisms and suggest novel therapeutic targets.

The brain has an intricate vascular network that satisfies the high energetic and metabolic demands of neuronal function through high levels and tight regulation of blood flow [21,22]. In the cortex, the main cerebral arteries on the brain surface branch into penetrating arterioles that plunge into the brain and feed the capillary network. The close proximity of capillaries to other types of cells such as neurons, astrocytes, microglia, and pericytes makes capillaries particularly important for the exchange of nutrients and oxygen to ensure proper brain function [23]. Previous studies have identified capillary abnormalities in AD patients [24–26] and mouse models of AD, such as focal reductions in capillary diameters due to pericyte constriction [8,27,28] and neutrophil arrest in capillary segments [29–32], which contribute to reduced CBF. In the APP/PS1 mouse model of AD, about 2% of cortical capillaries are stalled with an arrested neutrophil whereas only ~0.4% are stalled in wild-type (WT) mice. Reducing the number of stalls with an antibody against a neutrophil-specific surface protein (Ly6G) improved overall penetrating arteriole flow by ~32%, and cortical perfusion by ~18%. This flow increase was correlated with a rapid improvement in performance on spatial and working memory tasks [29], even in APP/PS1 mice at late stages of disease progression [33]. If this same mechanism contributes to CBF reduction in AD patients [34], and if the impact of improved CBF on cognitive function is as significant as in AD mouse models, decreasing the incidence of capillary stalling could represent a novel avenue for an effective therapy for AD. However, the molecular pathways driving increased capillary stalling still need to be understood.

Some of the molecular mediators of impaired neurovascular regulation have been uncovered in mouse models of AD. Briefly, amyloid-beta activates scavenger receptors in vascular associated cells [35] that trigger the activation of super-oxide-producing enzymes such as NADPH-oxidases, which are found with different catalytic subunits. The NOX2-containing NAPDH-oxidase isoform has been shown to be the most significant source of vascular reactive oxygen species (ROS) in mouse models of Aβ overexpression [36]. The ROS reacts with nitric oxide to form peroxynitrate, which causes DNA damage in endothelial cells, leading to activation of the DNA repair enzyme poly-ADP ribose polymerase (PARP-1) [37]. One downstream consequence of PARP-1 activation is the opening of TRPM2 channels in endothelial cells [38], leading to large increases in intracellular Ca^2+^, and contributing to the endothelial dysfunction that impairs neurovascular regulation. This ROS/peroxynitrate-mediated damage to endothelial cells may also drive low-level chronic inflammation that leads to increased expression of adhesion receptors, such as vascular cellular adhesion molecule 1 (VCAM-1) and intercellular adhesion molecule 1 (ICAM-1), in the endothelial cells of some capillaries, a pathway that may contribute to the arrest of neutrophils in capillary segments.

Here, we test the hypothesis that the NOX2 oxidative stress pathway contributes to vascular inflammation and capillary stalling. We used *in vivo* two photon imaging, in combination with gene expression profiling in cerebral microvessels, to determine the effect of inhibiting NOX2, as well as the downstream players PARP-1 and peroxynitrate [38], on capillary stalling, CBF, vascular inflammation, gene expression, and cognitive decline in 10-11 months old APP/PS1 mouse model of AD.

## 2. MATERIALS AND METHODS

### 2.1 Animals

All animal procedures were approved by the Cornell Institution Animal Care and Use Committee (IACUC, protocol number: 2015-0029) and were performed under the guidance of the Cornell Center for Animal Resources and Education (CARE). We used APP/PS1 transgenic mice (B6.Cg-Tg (APPswe, PSEN1dE9) 85Dbo/J; MMRRC_034832-JAX, The Jackson Laboratory) as a mouse model of AD, with wild-type (WT) littermates as controls. The APP/PS1 double transgenic mouse model expresses a chimeric mouse/human amyloid precursor protein (APP; Mo/HuAPP695swe) and contains a mutant human transgene for presenilin 1 (PS1; PSE-dE9). WT littermates served as controls. Animals were of both sexes and ranged from 10 to 11 months in age at the beginning of the study.

### 2.2 Inhibition of NOX2, PARP-1, and peroxynitrate activity

Inhibition of NOX2 activity was accomplished with an i.p. injection of 10 mg/kg in saline of gp91-ds-tat peptide (AS-63818, AnaSpec Inc. Fremont, CA) every other day for one and two weeks. We used a scrambled peptide, sgp91-ds-tat, as a control (AS-63821). Dosage and administration protocols were adapted from [39,40]. Inhibition of PARP-1 was accomplished by a 10-mg/kg i.p injection of 3-aminobenzamide in saline (PARP-1 inhibitor) every other day for a week, as described in [41]. The peroxynitrate decomposition catalyst FeTPPS (Millipore Sigma) was used to reduce peroxynitrate concentrations, *in vivo*. A dose of 30 mg/kg diluted in sterile saline was applied i.p. one hour and 24 hours (as described in [42]) after baseline imaging. We used saline vehicle injections as controls for the PARP-1 inhibitor and the FeTPPS.

### 2.3 ROS detection in brain slices

A first cohort of animals was used for ROS detection in the brain. As described in [43], dihydroethidine (DHE; Molecular probes) stock of 1.25 mg/mL was made fresh in anhydrous dymethilsufoxide (DMSO) and kept in the dark. Before injection, DHE stock was added to an equal volume of sterile saline and mixed. Mice were injected i.p. with 50 mg/kg of DHE (2 µM). To label blood vessels, a 0.5 mg/mL dose of FITC-Tomato Lectin (Thermofisher) was administered retro-orbitally. After one hour, mice were euthanized by a lethal injection of pentobarbital (50 mg/kg), brains were extracted, flash frozen and kept at -80 C° until the time of slicing. Brains were then sectioned at 30-µm thickness using a cryotome. Mounted tissue slices were imaged on a confocal microscope (Zeiss LSM710) using a laser excitation wavelength of 568 nm and an emission filter centered at 590 nm for DHE, and 480 nm excitation and an emission filter centered at 517 nm for FITC. For each tissue slice, four fields were randomly selected and imaged using a 25X oil immersion objective. Images were thresholded by a blinded experimenter and integrated density was measured per image using ImageJ and averaged results per mouse were reported.

### 2.4 Real time qPCR analysis

Brains from a second cohort of animals were used for RT-qPCR. Cortex and hippocampus of half brains were separated and snap frozen in liquid nitrogen. A ceramic mortar cooled with liquid nitrogen was used to grind frozen brain pieces until they became powder. The powder was then dissolved with RLT buffer from RNAeasy Mini Kit (Qiagen) and RNA was extracted using the manufacturer’s protocol. RNA integrity was measured using a bioanalyzer and RNA quantity was quantified using Qubit (ThermoFisher). cDNA synthesis was accomplished with SuperScript IV VILO Master Mix with ezDNAse Enzyme (ThermoFisher) and RT-qPCR was performed using Power SYBR Green Master Mix (ThermoFisher). Primers were acquired from OriGene and the following sequences for NOX2 (Mus musculus *Cybb*) were used: Forward 5’-3’: TGGCGATCTCAGCAAAAGGTGG, Reverse 5’-3’: GTACTGTCCCACCTCCATCTTG. Expression was normalized using the housekeeping gene beta-actin with the following sequences: Forward 5’-3’ CATTGCTGACAGGATGCAGAAGG, Reverse 5’-3’ TGCTGGAAGGTGGACAGTGAGG. The fold expression relative to beta-actin was determined by the ΔΔCt method.

### 2.5 Animal surgery and in vivo imaging with two-photon microscopy for blood flow measurements and capillary stalling

A third cohort of animals was used for *in vivo* imaging. The surgery and *in vivo* two-photon microscopy were performed as previously described [29]. In brief, a 8-mm diameter craniotomy was prepared over the parietal cortex and covered by gluing a glass coverslip to the skull. Animals rested for at least 4 weeks before imaging to minimize the impact of post-surgical inflammation. During the imaging session, mice were placed on a custom stereotactic frame and were anesthetized using ~1.5% isoflurane in 100% oxygen, making small adjustments to maintain the respiratory rate at ~1 Hz. The mouse was kept at 37 °C with a feedback-controlled heating pad (40-90-8D DC, FHC). After being anesthetized, mice received a subcutaneous injection of atropine (0.005 mg/100 g mouse weight; 54925-063-10, Med Pharmex) to prevent lung secretions and 5% glucose in saline (1 mL/100 g mouse weight) every hour during the experiment to prevent dehydration. For imaging, white blood cells and platelets were labeled with a retro-orbital injection of Rhodamine 6G (0.1 mL, 1 mg/mL in 0.9% saline, Acros Organics, Pure). Leukocytes were distinguished from blood platelets with a retro-orbital injection of Hoechst 33342 (50 μL, 4.8 mg/ml in 0.9% saline, Thermo Fisher Scientific). Texas Red dextran (50 μL, 2.5%, molecular weight = 70,000 kDa, Thermo Fisher Scientific) in saline was injected retro-orbitally just before imaging to label the vasculature. Mice were imaged on a custom-made two-photon excited fluorescence (2PEF) microscope, using an 830-nm excitation wavelength (Vision II, Coherent). Emission was detected on three photomultiplier tubes through the following emission filters: 417/60 nm for Hoechst, 550/49 nm for Rhodamine 6G, and 641/75 nm for Texas Red. We collected three-dimensional volumetric image stacks of ~300 × 300 × 300 µm^3^ to visualize arterioles, venules, and capillaries. Penetrating arterioles were identified based on their connectivity to surface arterioles and by determining their flow direction using the motion of unlabeled red blood cells in the fluorescently-labeled blood plasma. In each mouse, we identified 6-8 penetrating arterioles and took small 3D image stacks to determine the diameter as well as line scans along the centerline of the vessel to determine red blood cell flow speed, as described previously [44]. Capillary stalls were identified manually, and their cellular composition was identified with Hoescht and Rhodamine 6G. To extract the number of capillary segments in each image stack, we used a convolutional neural network, DeepVess [45], that segments 3D vessels within volumetric 2PEF images. Imaging timepoints included baseline imaging, 24 hours, and 7 days post-treatment.

### 2.6 Behavioral assays

To evaluate working and spatial short-term memory function, we used the Y-maze test and object displacement (OD) test. Behavioral analyses were performed at baseline, and after 1 week and 2 weeks of NOX2 inhibition. The Y-maze was made of light grey plastic and consisted of three arms at 120°. Each arm was 6-cm wide and 36-cm long and had 12.5-cm high walls. A mouse was placed in the Y-maze and allowed to explore for 6 min. The maze was cleaned with 70% ethanol after each mouse. Mouse behavior was monitored, recorded with a CCD camera, and analyzed manually by a blinded experimenter. A mouse was considered to have entered an arm if the whole body (except for the tail) entered the arm and to have exited if the whole body (except for the tail) exited the arm. An alternating trial was counted if an animal consecutively entered three different arms. Because the maximum number of trials is the total number of arm entries minus 2, the spontaneous alternation score was calculated as (number of alternating triads) / (total number of arm entries − 2).

For the OD task, all objects used were first validated in a separate cohort of mice to ensure that no intrinsic preference or aversion was observed, on average, and animals explored all objects similarly. Objects and arena were cleaned with 70% ethanol before and after each trial. Exploration time for the objects was defined as any time when there was physical contact with an object (whisking, sniffing or touching) or when the animal was oriented toward the object and the head was within 2 cm of it. In trial 1, mice were allowed to explore two identical objects for 10 min in the arena and then returned to their home cage for 60 min. Mice were then returned to the testing arena for 3 min with one object moved to a novel location (trial 2). We ensured that the change of placement altered both the intrinsic relationship between objects and the position relative to internal visual cues. At subsequent time points, new object positions and new pairs of objects (from the validated pool of objects) were used to maintain animal interest. In addition to using the tracking software (Viewer III; Biobserve, Bonn, Germany) to determine the object exploration times, the time spent at each object was manually scored by a experimenter who was blinded to the genotype and treatment. The preference score (%) for object displacement tasks was calculated, from the data in trial 2, as ([exploration time of the novel object] / [exploration time of both objects]) x 100. Automated tracking and manual scoring yielded similar results across groups, so we report the automated tracking results.

### 2.7 Immunohistochemistry

Animals from the third cohort were euthanized by lethal injection of pentobarbital (5 mg/100 g) and perfused transcardially with 1X PBS after imaging. Brains were quickly extracted and divided along the centerline. One half was immersed in 4% paraformaldehyde in phosphate buffered saline (PBS) for later histological analysis and the other half was snap frozen in liquid nitrogen. As described in [46], brain halves were embedded in OCT medium and positioned on a cryotome to perform 30-µm sagittal cuts. Slices were then mounted on histology slides, washed 5 times with 1X PBS and incubated for 1 hour with blocking solution (Goat serum, 10% TritonX, 1X PBS). Primary antibodies for GFAP (anti-chicken, Rockland) and IBA1 (anti-rabbit, WAKO chemicals), were applied in a 1:500 dilution factor with blocking solution, and slides were incubated at 4°C overnight. On the next day, slides were washed 3 times with 1X PBS, and the corresponding secondary antibodies (Alexa Fluor 488, or 594) were applied in a 1:500 dilution factor and incubated for 2 h at room temperature. Slides were washed a final time 3 times with 1X PBS and stained with a solution of methoxy-X04 (5 mg/mL in 10% DMSO, 45% propylene glycol, 45% 1X PBS [47]) and incubated for 15 minutes at room temperature. Then, slides were washed 2 times with 80% ethanol and allowed to dry, mounted with Prolong Gold (Thermofisher) mounting medium to avoid photobleaching, and imaged with a Zeiss 710 Confocal Microscope equipped with a 25X oil immersion objective. Three images of the hippocampus and cortex were taken per animal, and a blinded experimenter calculated the % area above a manually-set threshold using ImageJ.

### 2.8 ELISA assays

The other brain halves (frozen) were homogenized in 1 mL PBS containing complete protease inhibitor (Roche Applied Science) and 1 mM AEBSF (Sigma) using a Dounce homogenizer. The homogenates were then sonicated and centrifuged at 14,000 g for 30 min at 4° C. The supernatant (PBS-soluble fraction) was removed and stored at −80° C. The pellet was re-dissolved in 0.5 ml 70% formic acid, sonicated, and centrifuged at 14,000 g for 30 min at 4° C, and the supernatant was removed and neutralized using 1M Tris buffer at pH 11. Protein concentration was measured in the PBS soluble fraction and the formic acid soluble fraction using the Pierce BCA Protein Assay (Thermo Fischer Scientific). The PBS soluble fraction extracts were diluted 1:5. Formic acid extracts were diluted 1:1 after neutralization. These brain extracts were analyzed by sandwich ELISA for Aβ1-40 and Aβ1-42 using commercial ELISA kits and following the manufacturer’s protocol (Aβ1-40: KHB348 and Aβ1-42: KHB3441, Thermo Fisher Scientific). The Aβ concentration was calculated by comparing the sample absorbance with the absorbance of known concentrations of synthetic Aβ1–40 and Aβ1–42 standards on the same plate. Data was acquired with a Synergy HT plate reader (BioTek) and analyzed using Gen5 software (BioTek) and Prism (GraphPad) [46].

### 2.9 Extraction of RNA from cerebral microvessels

A cohort of mice (10-11 months, n = 8 per group, both APP/PS1 and WT genotypes) were used for RNA sequencing of cerebral microvessels. Briefly, animals were treated with gp-91-ds-tat or its scrambled control (sgp-91-ds-tat) for two weeks. After treatment, they were euthanized with an intraperitoneal injection of pentobarbital (100 µL) followed by a cervical dislocation, and brains were immediately extracted and were transferred to cold MCDB 131 medium (Thermo Fisher Scientific, 10372019). The subsequent steps are explained in more detail in [48]. In brief, procedures were conducted in a cold room to minimize cell activation. The cortical tissue was homogenized with MCDB131 medium with 0.5% fatty acid free BSA (Millipore Sigma, 126609) using a Dounce homogenizer. The homogenate was centrifuged at 2000 g for 5 minutes at 4°C. The pellet was suspended in 15% dextran (molecular weight ∼70 kDa, Millipore Sigma, 31390) in PBS and centrifuged at 10000 g for 15 minutes at 4°C. The pellet was resuspended in MCDB131 with 0.5% fatty acid free BSA and centrifuged at 2000 g for 10 min at 4°C. The final pellet contained the microvessels. Samples were stored in -80ºC until further processing. Isolated cerebral microvessels were homogenized in appropriate amount of RLT lysis buffer to lyse the multicellular structures. Lysed samples were then loaded onto a column-based shredder (QIAshredder, Qiagen, Germany) and centrifuged (8000 g for 2 min at room temperature) to elute the homogenized lysate. Thereafter, the lysate was processed using total RNeasy mini kit (Qiagen, Germany). The lysate was resuspended in a 1:1 volume of 70% ethanol and loaded onto an extraction column and centrifuged (8000g for 2 min) at room temperature. DNase-I digestion was performed on-column for samples with 1U of TURBO-DNase (Invitrogen, ThermoFisher Scientific) and incubated for 15 min at room temperature. Then columns were washed with appropriate amount of wash buffer and finally total RNA was eluted using 20-25 µl of nuclease free water.

### 2.10 RNA-sequencing

RNA integrity and other quality control measures were performed along with ribosomal RNA depletion before cDNA library preparation (Genomics core facility, Weill Cornell Medicine). cDNA libraries were constructed using the Illumina TruSeq Stranded mRNA Library Prep kit and were sequenced in a single lane with pair-end 101 bps on Illumina HiSeq4000 instrument. The raw sequencing reads in BCL format were processed through bcl2fastq 2.20 (Illumina) for FASTQ conversion and demultiplexing. Cutadapt [49] was used to trim low quality bases and adapters. STAR (Version 2.5.2) [50] and Cufflinks (Version 2.1.1) were used to align raw sequencing reads to the mouse GRCm38 reference genome. The abundance of transcripts was measured with Cufflinks in fragments per kilo base of transcript per million mapped fragments (FPKM). Gene expression profiles were constructed for differential expression, principle component analyses and MA plots (log fold change vs mean of normalized counts) with the DESeq2 package [51]. For differential expression analysis, pairwise comparisons between two or more groups using parametric tests where read-counts follow a negative binomial distribution with a gene-specific dispersion parameter were used. Corrected p-values were calculated based on the Benjamini-Hochberg method to adjust for multiple comparisons. The following comparisons were performed for our data: AD vs WT and treated vs non-treated in all the groups. A sample-to-sample distance heatmap, volcano plots (log p-value versus log fold change), and principal component analysis (PCA) were done using iDEP [52]. Gene set enrichment analysis GSEA [53] was performed with our dataset to look for biological pathways enriched between groups. The normalized matrix of counts from DESeq2 was inputted onto GSEA, along with the sample metadata, and the program was run for the gene ontology category of biological process (GOBP). Gene sets that were positively or negatively enriched in our dataset were visualized as a network using Cytoscape with the EnrichmentMap and AutoAnnotate [54] plugin. Raw FASTQ datasets are publicly available in the Gene Expression Omnibus (GEO) with the accession number GSE224394.

### 2.11 In vivo immunofluorescence imaging of vascular adhesion molecules in brain vessels

A fifth cohort of animals was used for *in vivo* imaging of endothelial inflammatory receptors using multiphoton microscopy. As described in [55], the following antibodies were used for imaging: BV421 Hamster Anti-Mouse CD54 (ICAM-1) Clone 3E2 (BD Biosciences), and BV421 Rat Anti-Mouse CD106 (VCAM-1) Clone 429 (BD Biosciences). For *in vivo* imaging, a dose of 0.5 mg/kg was injected retro-orbitally into the vasculature, and imaging was performed 24 hours post-injection to maximize antibody binding and ensure clearance of unbound antibodies from the vasculature. A dose of Texas Red (50 μl, 2.5%, molecular weight = 70,000 kDa, Thermo Fisher Scientific) or FITC (50 μl, 2.5%, molecular weight = 70,000 kDa, Thermo Fisher Scientific) in saline was applied to label the vasculature. To maintain consistent labeling and minimize variation between animals, imaging parameters were kept the same through multiple imaging sessions, and we paid special attention to keeping the PMT gain, laser power, and wavelength (830 nm) the same between sessions. For the image analysis, an experimenter blinded to the animal phenotypes manually screened the composite 3D stacks and segmented the contour of 3-4 penetrating arterioles, capillaries, and venules per stack with the freehand tool in ImageJ. We acquired 4-6 stacks per animal, resulting in an average of 12-24 vessels measured in each animal. After the manual segmentation, the raw integrated density, mean intensity, and area in each fluorescent channel was measured, and the same measurements were taken in a region at the corner of the image for background measurements. The location in the stack in pixels (X,Y,Z) of each vessel was registered along with the fluorescence measurements. Background values were subtracted from the raw integrated density on each channel, and the resulting value was then divided by the intensity of the vascular label to normalize the data for depth-dependent loss of signal. Datapoints were pooled for each animal and represented as normalized intensity.

### 2.13 Staining of human cortical tissue

Human brain tissue was provided by the NIH NeuroBioBank, the Harvard Brain Tissue Resource Center, Sepulveda Research Corporation, the Mount Sinai/JJ Peters VA Medical Center NIH Brain and Tissue Repository and the University of Maryland Brain and Tissue Bank. Samples and their correspondent brain region, donor, age, and sex are located in Supplementary Table 1. Immunohistochemistry of human brain tissue followed the protocol in Ref. [56]. Briefly, *post-mortem* brain tissue fixed in 4% PFA was embedded in OCT adhesive compound, frozen, and sectioned using a cryostat into 40 µm thick slices. Slices were mounted on glass slides and washed three times with 1X PBS. To block endogenous peroxidases, the tissue was incubated for 20 min in 50% methanol with 1% H2O2. Sections were then washed and incubated for 30 min with 0.3 % Sudan black solution in 70% ethanol to reduce background autofluorescence. Slides were then washed in 1X PBS for 5 min, and stained with VECTASTAIN Elite ABC Universal Kit Peroxidase Horse anti-mouse/Rabbit IgG (Vector laboratories) using the primary antibody mouse anti-human g91-phox (Santa Cruz Biotechnologies, sc-130543) with a 1:100 dilution. Non-specific secondary labeling was assessed by omitting the primary antibody.

### 2.14 Statistical analysis

All analysis, except the RNAseq work and linear mixed effect models, was conducted on Prism8 (GraphPad). To analyze statistical differences between groups, data was first tested for normality with a D’Agostino-Pearson’s normality test. If data was normal, a one-way ANOVA was used in datasets with more than two comparison groups, or a t-test for paired or unpaired datasets. In the case of non-normal distribution, a Kruskal Wallis or Mann-Whitney test was used. In the cases where repeated measurements were performed across animals and timepoints, a linear mixed effect model was used to describe interactions between fixed and random effects, and to control for inter-subject variability. These analyses were performed in R with the lme4 package. Some logarithmic transformations of skewed data were performed and are indicated in the respective figure. P-values less than 0.05 were considered statistically significant while for values between 0.05 and 0.1 we reported a “trend” in the data. The following indicators of significance are used in the figures: * *P* < 0.05, ** *P* < 0.01, *** *P* < 0.001, **** *P* < 0.0001. In box plots, red bars in graphs indicate mean and black bars indicate median, and the box spans the 25^th^ to 75^th^ percentile of the data. Whiskers span from the lowest data point within 1.5 times the interquartile range (75^th^ -25^th^ percent; IQR) of the lower quartile of the data to the highest data within 1.5 times the IQR of the highest quartile of the data.

## 3. RESULTS

### 3.1 NOX2 inhibition reduced ROS generation as well as NOX2 mRNA expression by two weeks post treatment

We studied changes in CBF and capillary stalling using *in* vivo 2PEF imaging in craniotomized APP/PS1 and WT mice after treatment with a small peptide inhibitor of NOX2, gp91-ds-tat (10 mg/kg in saline), as compared to treatment with a scrambled control peptide, sgp91-ds-tat (Fig 1A). Animals were then euthanized for post-mortem tissue analysis.

**Figure 1.**
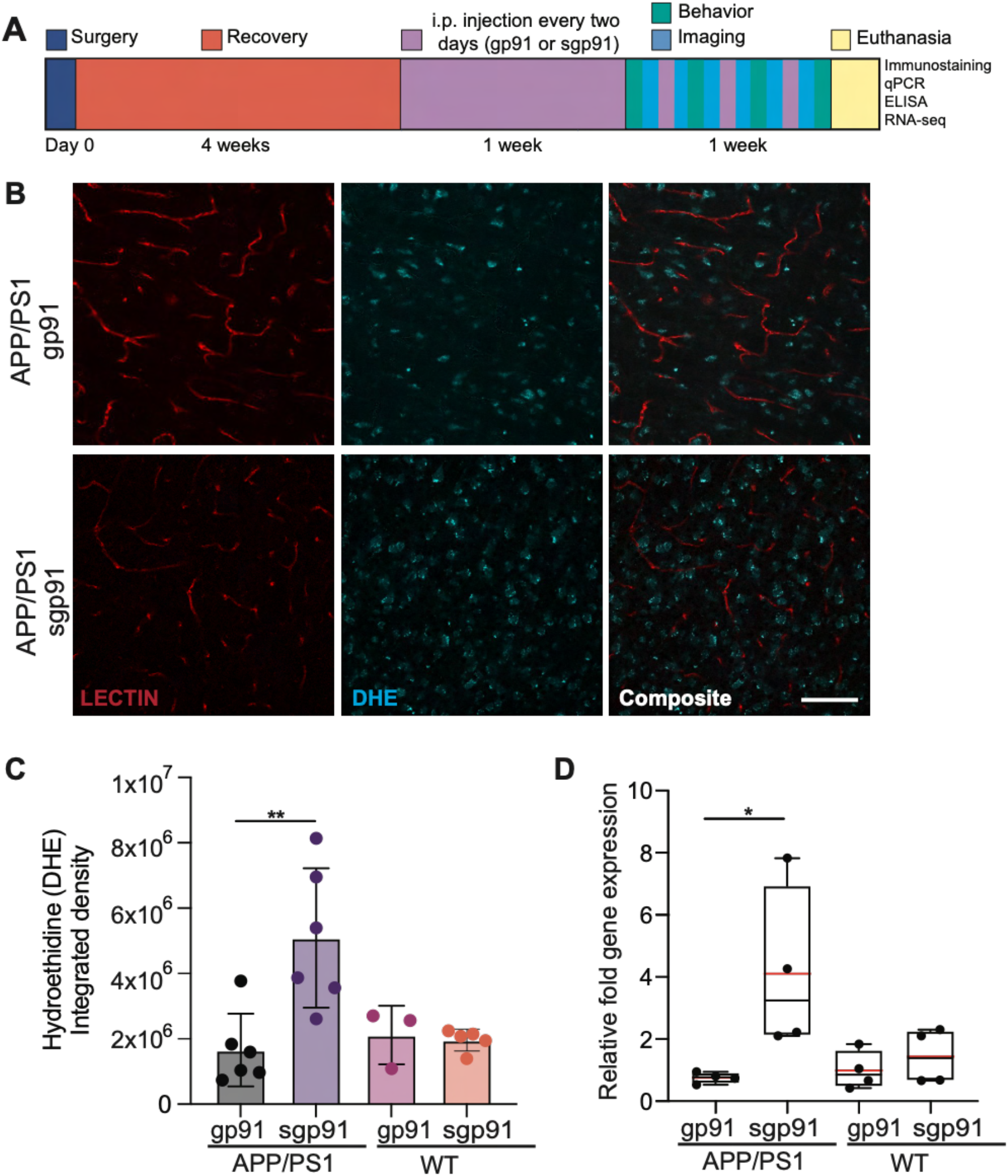
Treatment with the NOX2 inhibitor gp91 decreases the generation of reactive oxygen species and NOX2 mRNA levels in the neocortex of APP/PS1 mice after two weeks of treatment. **A)** Experimental design timeline. Following euthanasia, brains were extracted and analyzed for immunostaining, NOX2 mRNA, RNA seq, or protein extraction. **B)** Z-projections of confocal images of cortical brain sections stained with DHE and lectin for blood vessels. **C)** Integrated density measurements of DHE staining of APP/PS1 and WT mice treated with NOX2 inhibitor (gp91) or scrambled control peptide (sgp91) (WT gp91 n=3, WT sgp91 n=5, APP/PS1 gp91 n=6, APP/PS1 sgp91 n= 6; one-way ANOVA with Tukey post-hoc comparison). **D)** Relative fold NOX2 mRNA expression in treated and control mice assessed with RT qPCR (WT gp91 n=4, WT sgp91 n=4, APP/PS1 gp91 n=4, APP/PS1 sgp91 n= 4; one-way ANOVA, with Tukey post-hoc comparison). *p < 0.05, ** p < 0.01

To determine whether NOX2 inhibition was achieved in the brain of APP/PS1 mice, we quantified the production of ROS with hydroethidine staining (DHE) and the relative expression of *Nox2* in the cortex with RT-qPCR after two weeks of treatment. We detected a decrease in the amount of ROS accumulated in brain tissue (Fig 1B and 1C) in addition to a significant decrease in the relative fold mRNA expression of *Nox2* in APP/PS1 mice (Fig 1D), as compared to scrambled peptide control treatment. No differences were seen in WT animals between NOX2 inhibition and scrambled treatment.

### 3.2 NOX2 inhibition reduced capillary stalling in APP/PS1 mice after one week of treatment

From two-photon images of fluorescently labeled cortical vasculature (Fig. 2A), capillaries were scored as stalled or flowing based on the apparent motion of red blood cells over time (Fig 2B). Consistent with our previous work [29,30], we found a ~5X elevation in the incidence of stalled capillaries in APP/PS1 as compared to WT mice. Treatment with the NOX2 inhibitor had no impact after 1 hour on capillary stalling, but led to a 67% decrease in the incidence of capillary stalls after 1 week in APP/PS1 mice, while the scrambled peptide had no effect (Fig. 2C). We also observed a trend toward decreased capillary stalling after treatment with the peroxynitrate decomposition catalyst FeTPPS (Sup Fig. 1A), as well as after inhibition of the DNA damage responding enzyme PARP-1 (Sup Fig. 2A). Using additional fluorescent labels (see Methods) we distinguished the cellular cause of the stalled capillaries (Fig. 2D) and found that the majority of stalls in APP/PS1 mice were caused by an arrested white blood cell. Although NOX2 inhibition reduced the incidence of stalls, it did not significantly change their cellular composition, i.e., the type of cell causing the stalls (Fig. 2E).

**Figure 2.**
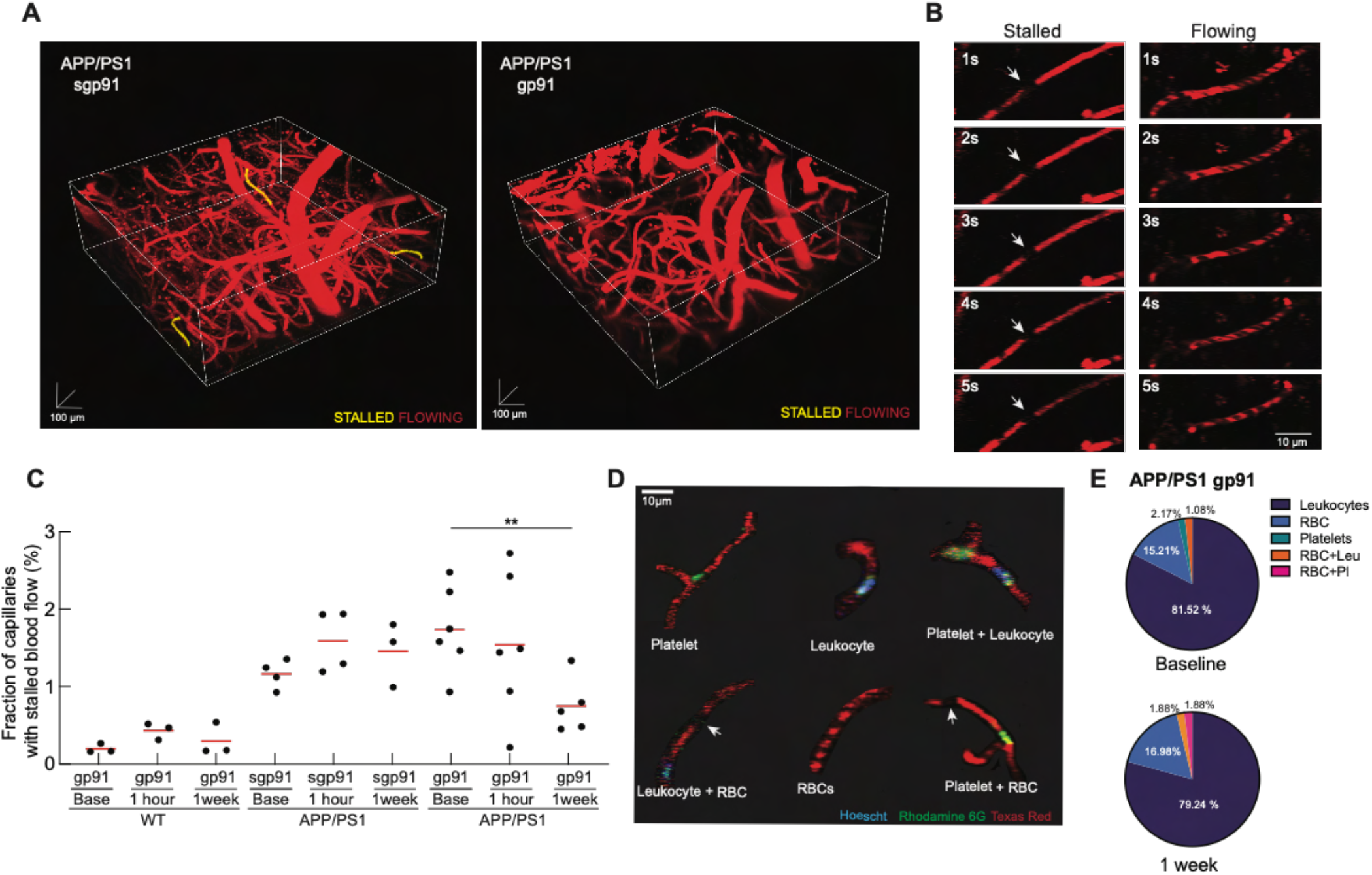
NOX2 inhibition decreased the incidence of capillary stalls after 1 week of treatment. A) Representative 3D image stacks of APP/PS1 mice treated with the NOX2 inhibitor gp91 or scrambled control sgp91. B) Representative 2PEF sequences of stalled and flowing capillaries over time. C) Fraction of capillaries with stalled blood flow, one hour and one week after treatment with NOX2 inhibitor or scrambled control for all groups. D) Types of cells composing capillary stalls (Hoescht + Rhodamine 6G: leukocytes; Rhodamine 6G: platelets; Dark spots: RBCs). E) Pie charts representing the percentage of cell types responsible for capillary stalling in APP/PS1 animals treated with NOX2 inhibitor (gp91) at baseline and 1 week post treatment. RBC = red blood cells, Leu = leukocytes, Pl = platelets. (WT gp91 n=5, APP/PS1 gp91 n=6, APP/PS1 sgp91 n= 6; one-way ANOVA with Tukey post-hoc comparison). **p < 0.01.

### 3.3 NOX2 inhibition increased blood flow in penetrating arterioles in APP/PS1 mice after one week of treatment

Because NOX2 inhibition reduces capillary stalling in APP/PS1 mice, we sought to determine if there were changes in CBF in NOX2-treated animals. Previous studies demonstrated that NOX2 [38,57,58] and PARP-1 [38] inhibition improves blood flow attenuation induced by neocortical superfusion of amyloid-beta peptide in WT mice [58]. These studies utilized laser doppler to estimate blood flow and were unable to resolve the cellular mechanism underlying the Aβ triggered blood flow reduction nor the rescue with NOX2 or PARP-1 inhibition. Using linescans to quantify RBC flow speed (Fig. 3A), we found that NOX2 significantly increased volumetric blood flow (Fig 3B) and RBC flow speed (Fig. 3C), but had no effect in vessel diameter (Fig. 3D) of cortical penetrating arterioles, as compared to the scrambled peptide and to WT controls. We observed similar effects after treatment with FeTPPS (Sup Fig. 1B) and PARP-1 inhibition (Sup Fig. 2B – D).

**Figure 3.**
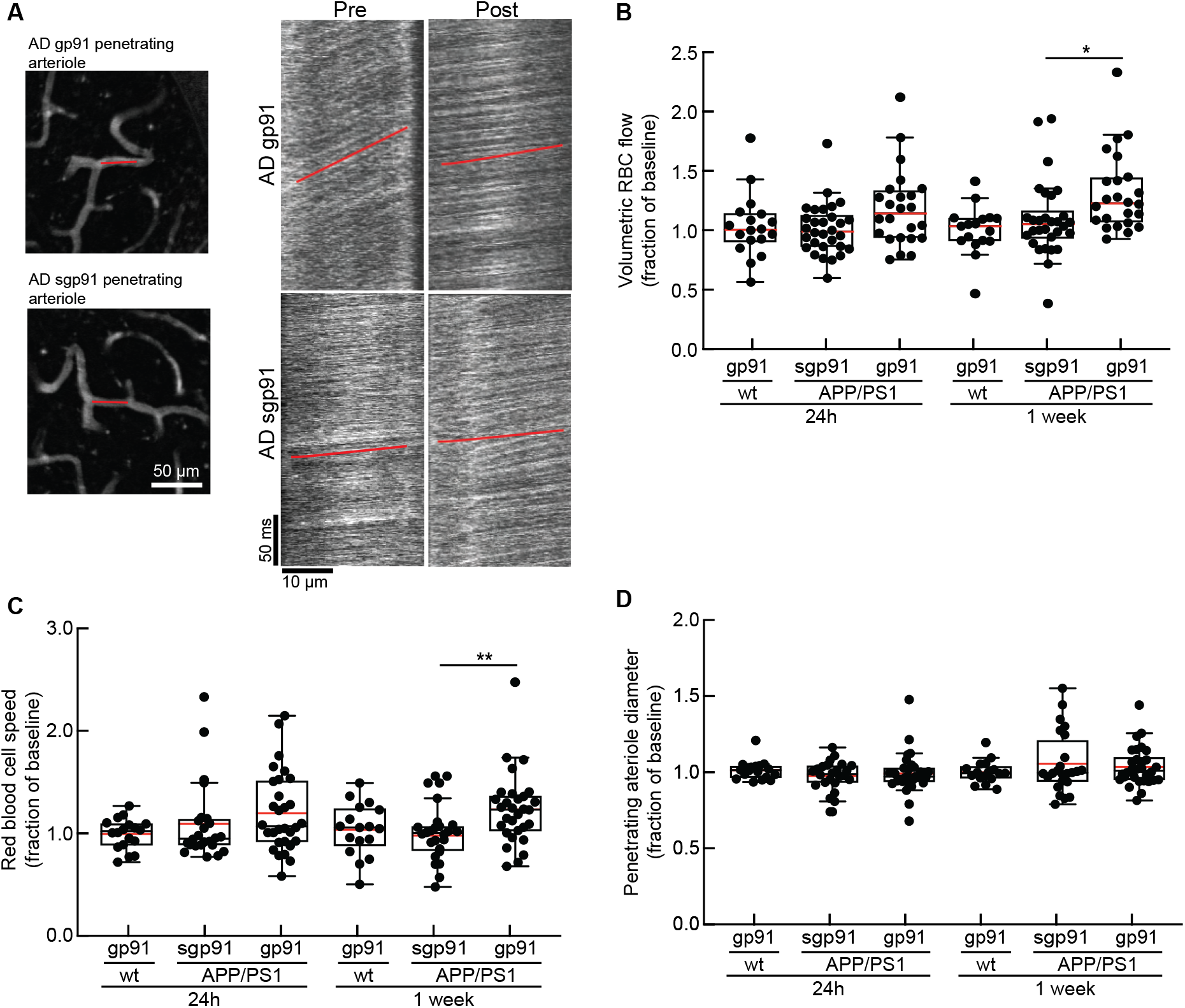
NOX2 inhibition improves blood flow in penetrating arterioles in APP/PS1 mice after one week of treatment. A) Representative examples of linescans from penetrating arterioles in APP/PS1 mice treated with NOX2 inhibitor gp91 or scrambled control sgp91. The red lines represent the scanning direction along the vessel centerline. The slope (red line) of the black stripes in the linescan images is used to extract the RBC speed. A steeper slope corresponds to slower flow. B) Volumetric blood flow, C) red blood cell speed and D) diameter from penetrating arterioles of APP/PS1 mice and WT mice treated with gp91 or scrambled control sgp91. Significance was tested with a linear mixed effect model (estimate of fixed effects of control vs treatment: -0.02, *p < 0.05, **p < 0.01). n = 6 penetrating arterioles per mouse, with animals in groups: WT gp91 n=3, APP/PS1 gp91 n=4, APP/PS1 sgp91 n= 5.

### 3.4 NOX2 inhibition improves short-term and spatial memory in APP/PS1 mice after two weeks of treatment

Previous studies showed that NOX2 inhibition improves short term memory in mice overexpressing APP with Swedish mutations [36]. Here we evaluate whether memory function, as assayed by the object displacement (Fig. 4A) and the Y-maze spontaneous alternation (Fig. 4B) tests, is improved in APP/PS1 mice after NOX2 inhibition, in parallel with the reduction in capillary stalls and the increase in CBF.

**Figure 4.**
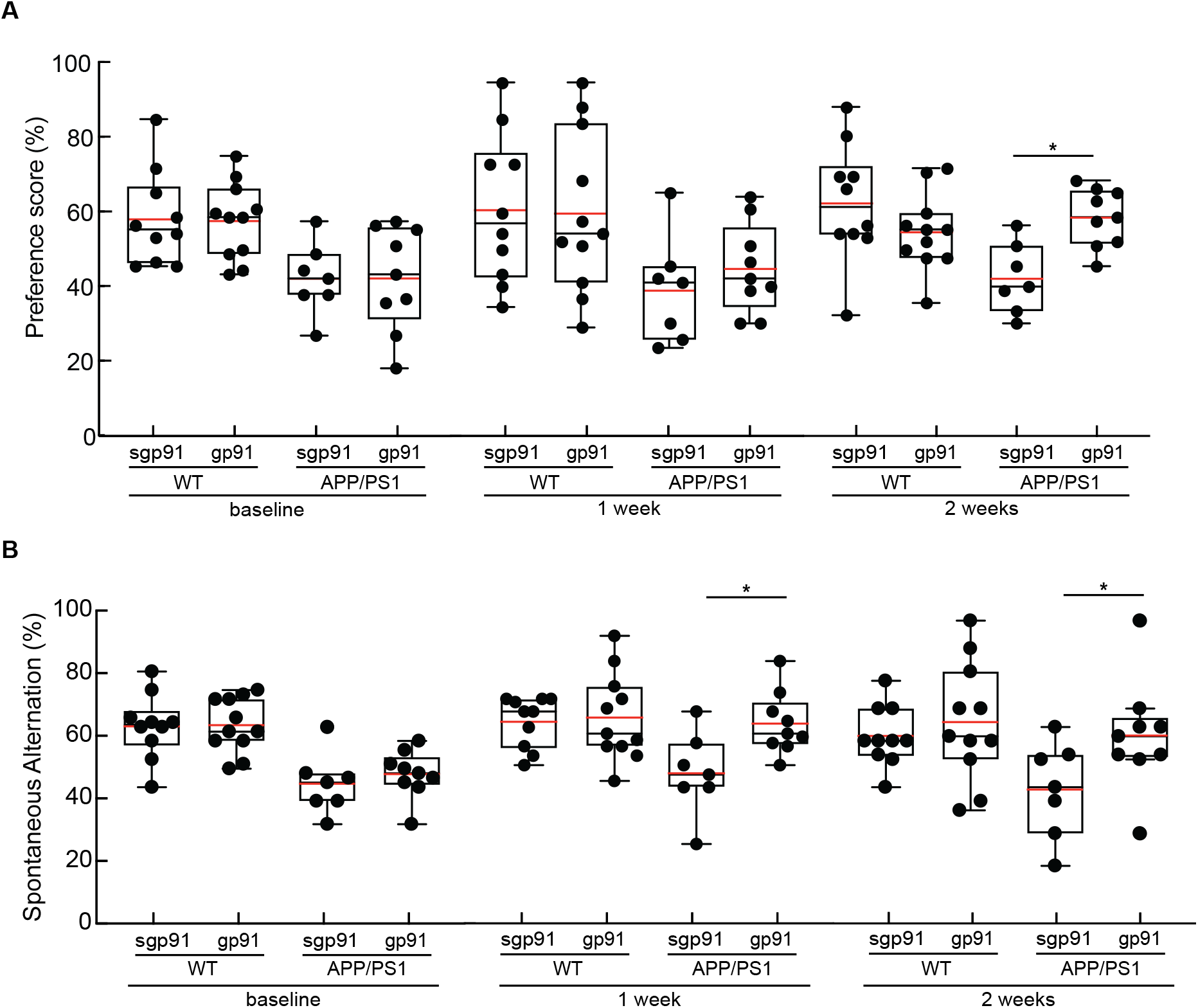
NOX2 inhibition improves short-term and spatial memory functions in APP/PS1 mice after two weeks of treatment. **A)** OD preference score and B) Y-maze spontaneous alternation score for APP/PS1 and WT mice treated with gp91 or scrambled control sgp91 at baseline, and after 1 week and 2 weeks of treatment every other day. Animal numbers: APP/PS1 sgp1 n=7, APP/PS1 gp91 n=9, WT gp91 n= 11, WT sgp91 n= 10. Significance was tested with a mixed-effects model and Tukey’s multiple comparisons test (fixed effect of treatment (gp91 vs sgp91) and group (AD vs WT), * p < 0.05). Fixed effect of timepoint was not statistically significant (p = 0.48).

We performed behavioral testing in APP/PS1 mice after one and two weeks of NOX2 inhibition. Although we saw a decrease in the number of capillary stalls and increase in CBF after one week of NOX2 inhibition, we observed only a slight trend toward improved performance on the OR test, and a significant improvement in the Y-maze at one week, with a larger and statistically significant improvement evident at two weeks for both tests. The number of arm entries and time spent at the replaced object remained unchanged between groups, suggesting continued engagement with the memory task across the three timepoints (Sup Fig. 3A, 3B). These results indicate that NOX2 inhibition also contributes to an improvement in short-term and spatial memory functions in APP/PS1 mice.

### 3.5 NOX2 inhibition ammeliorates neuroinflammation independent of amyloid-beta in APP/PS1 mice

Neuroinflammation and amyloid-beta accumulation are closely associated with vascular oxidative stress and blood flow alterations [59]. Thus, we examined whether the reduction in capillary stalls and rescue in arteriole blood flow observed in APP/PS1 mice after NOX2 inhibition are accompanied by a reduction in neuroinflammation and amyloid-beta levels. First, we used microglia/macrophage and astrocytic density identified with IBA1 and GFAP staining, respectively, as markers of neuroinflammation, in cortical and hippocampal slices of mice treated with NOX2 inhibitor for two weeks (Fig. 5A). We found a significant decrease in GFAP expression in the hippocampus (Fig. 5C), but not in cortex (Fig. 5B), and a decrease in IBA 1 expression in the cortex (Fig. 5D) and hippocampus (Fig. 5E) in NOX2 inhibited APP/PS1 mice. To address if other antioxidants besides gp91-ds-tat help to reduce inflammation in the cortex, we validated our results using apocynin, a common non-specific NOX inhibitor [60]. We also found that apocynin reduces IBA1 positive cells in brain cortical tissue of APP/PS1 mice (Sup Fig 4A). We used Methoxy-X04 staining to quantify plaque load in cortical and hippocampal slices of the same mice. We found no differences in Methoxy-X04 staining between treated and untreated mice in either cortical (Fig. 5F) or hippocampal (Fig. 5G) slices, nor differences in the average density of plaques (Fig. 5H). We also did not detect changes in the soluble fraction of amyloid beta 40 (Sup Fig. 5A), amyloid beta 42 (Sup Fig. 5B), or insoluble amyloid beta 42 (Sup Fig 5C), consistent with previous studies [36]. Finally, we did not detect any changes in the average number of microglia (Fig. 5I) or astrocytes (Fig. 5J) surrounding amyloid plaques. These findings indicate that NOX2 inhibition ameliorates neuroinflammation, independent of amyloid-beta levels, in APP/PS1 mice.

**Figure 5.**
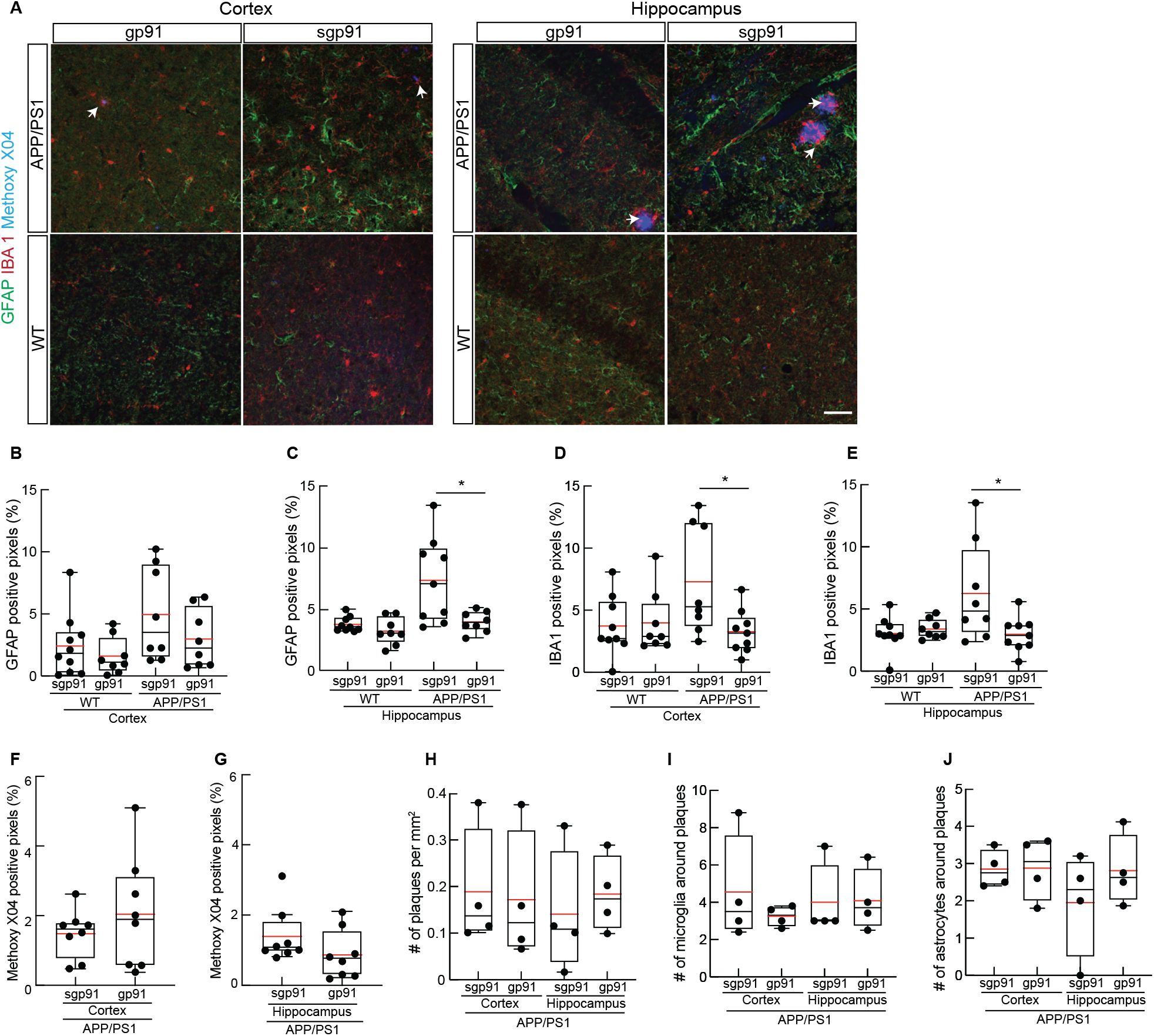
NOX2 inhibition ameliorates neuroinflammation independent of amyloid-beta levels in APP/PS1 mice. **A)** Representative confocal images of brain tissue sections from cortex and hippocampus of APP/PS1 mice treated with gp91 (left) or sgp91 (right), labeled with anti-GFAP (astrocytes; green), anti-IBA1 (microglia/macrophage; red) and Methoxy X04 (amyloid-beta plaques; blue, white arrowheads). **B)** Astrocytic density in cortex and **C)** hippocampus of APP/PS1 mice treated with gp91 or sgp91 (expressed as the percentage of GFAP-positive pixels). **D)** Microglial density in cortex and **E)** hippocampus of APP/PS1 mice with gp91 or sgp91 (expressed as the percentage of IBA1-positive pixels). Amyloid plaque load represented as the percentage of pixels that are Methoxy X04 positive in **F)** cortical and **G)** hippocampal brain slices. **H)** Number of plaques per unit area from APP/PS1 mice in cortex and hippocampus. **I)** Number of microglia and **J)** number of astrocytes surrounding plaques in APP/PS1 mice in cortex and hippocampus. Animal numbers: WT gp91 n=8, WT sgp91 n=9, APP/PS1 gp91 n=8, APP/PS1 sgp91 n=9. Significance was measured with a Kruskal Wallis test with Tukey post-hoc multiple comparisons correction (* p < 0.05). Scale bar = 50 µm.

### 3.6 RNA sequencing and gene set enrichment analysis (GSEA) of cerebral microvessels reveals differentially expressed genes involved in inflammation and leukocyte adhesion sensitive to NOX2 inhibition in APP/PS1 mice

To investigate the molecular mechanisms contributing to increased capillary stalls and the impact of NOX2 inhibition in APP/PS1 mice, we performed differential gene expression analysis using the transcriptome of cerebral microvessels extracted from APP/PS1 and WT mice. We used an approach to extract cerebral microvessels that minimizes changes in the inflammatory profile of endothelial and other vascular associated cells [48]. We then performed RNA sequencing on the cerebral microvessels extracted from WT and APP/PS1 mice, treated with the NOX2 inhibitor or the scrambled control (Fig. 6A). A sample-to-sample heatmap was used to visualize overall gene expression across all samples (Fig. 6B). We first compared gene expression between APP/PS1 and WT, and analyzed them using DESeq2 and GSEA. We found 29 upregulated genes and 0 downregulated genes (log2FC > 1, padj <0.1, Fig 6C). Among the more strongly upregulated genes, we found the cystatin F (*Cst7*), which is neutrophil-specific and commonly associated with cerebral amyloid angiopathy [61], the astrocyte marker *Gfap* indicative of glial activation and commonly upregulated in AD [62,63], *Trem2* as a risk gene and common inflammatory signature marker abundant in microglia, as well as *Itgax, Cd68* and *Ccl3*. These results are consistent with previous gene expression analyses of endothelial cells between WT and APP/PS1 mice [64]. Gene set enrichment analysis (GSA) revealed an upregulation of gene sets linked to immune response, including regulation of leukocyte activation, interferon-gamma production, tumor necrosis factor (TNF) production, among others (FDR < 0.075, Fig 6D). Next, we explored the differential response of NOX2 treatment (gp91) in comparison to control (sgp91). Contrary to our hypothesis, we did not observe any differentially expressed (DE) genes between NOX2 inhibited and scrambled control mice (Sup Fig. 6C) in either APP/PS1 or WT groups, as assessed with the DESeq2 pipeline (log2FC >1.5, padj < 0.1). We did, however, observe significant changes at a pathway level with GSEA (FDR < 0.075). By comparing APP/PS1 mice with NOX2 inhibitor (gp91) vs scrambled control (sgp91), GSEA revealed downregulation of several pathways involved in inflammation and immune response, such as interferon gamma production (*Arg1, Trem1, Il6, Il1b, Il10*), heterotypic cell adhesion (*Vcam1, Nrcam, Il1b, Itgad, Tnf*), cell mediated integrin expression (*Serpine 1, Itgam, Cxcl13*), leukocyte migration and chemotaxis (*Cxcl13, Ccr7, Ccr6, Cxcr2)*, and plasma adhesion molecules (*Cldn8, Cdhr5, Crtam, Cdh17*) (Fig. 6E). These results overall indicate that cortical microvessels in AD mice are susceptible to inflammation and oxidative stress, and that NOX2 inhibition is able to downregulate some of these pathways.

**Figure 6.**
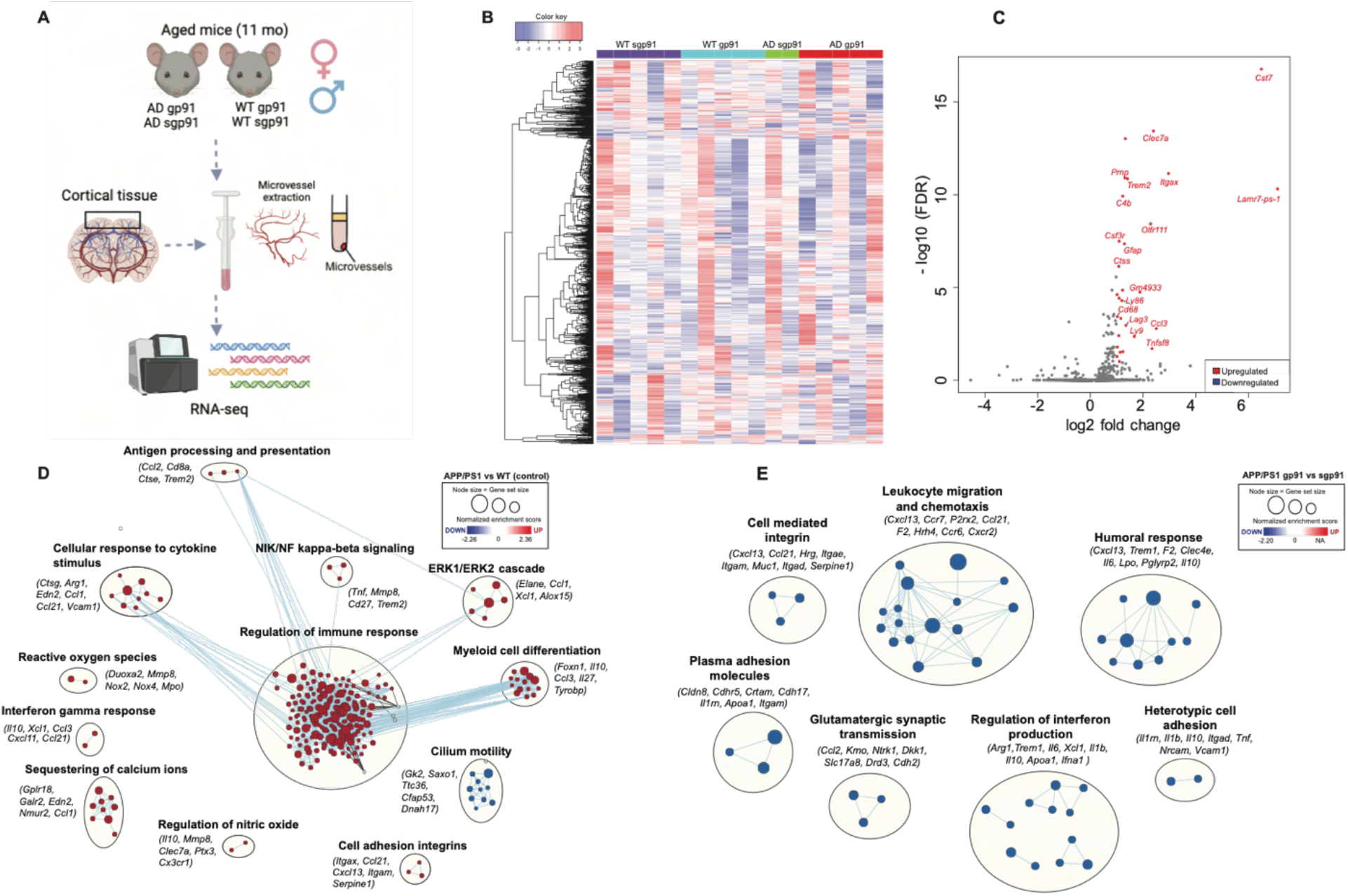
RNA sequencing and gene set enrichment analysis (GSEA) of cerebral microvessels reveals differentially expressed genes involved in inflammation and leukocyte adhesion sensitive to NOX2 inhibition in APP/PS1 mice. A) Experimental design and overall procedure of RNA-seq of microvessels. B) Unclustered heatmap of overall gene expression for all phenotypes (APP/PS1 vs WT) and treatments (NOX2 gp91 or scrambled control sgp91). C) Volcano plot of differentially expressed genes between APP/PS1 control and WT control (log2FC > 1.5, padj < 0.1). D) Enrichment network map of significantly up and down regulated gene ontology gene sets (FDR < 0.075) between APP/PS1 mice and WT with control treatment. E) Enrichment network map of significantly up and down regulated gene ontology sets (FDR < 0.075) between AD gp91 (NOX2 inhibited) and AD sgp91 (control). Node size indicates gene set size, similar gene sets that have overlapping genes are connected by edges (light blue lines). Node colors indicate the normalized enrichment score (NES). Animal numbers: WT sgp91 n=5, WT gp91 n=5, APP/PS1 sgp91 n=2, APP/PS1 gp91 n=5.

### 3.7 *In vivo* immunolabeling of adhesion receptors in blood vessels reveals an increase in the expression of VCAM-1 and ICAM-1 in APP/PS1 mice

Because capillary stalling contributes to CBF deficits in APP/PS1 mice that is corrected by NOX2 inhibition, we focused on gene sets upregulated in APP/PS1 mice (Fig 6D) that were downregulated with NOX2 inhibition (Fig 6E). Several gene sets that fit this pattern were related to cellular adhesion. To corroborate these findings, we used an *in vivo* labelling strategy [55] to compare the upregulation of endothelial inflammatory adhesion receptors, such as vascular cellular adhesion molecule -1 (VCAM-1) and intercellular adhesion molecule -1 (ICAM-1) throughout the vascular network. Considering that capillary stalls are unique *in vivo* phenomena, *in vivo* experiments are required to determine whether the expression of such adhesion molecules exists in stall-prone vs. flowing capillaries. Mice underwent treatment with NOX2 inhibitor for two weeks and then imaging was performed (Fig. 7A). We tested the antibody labelling specificity by comparing our imaging to the administration of fluorescently labelled isotype control antibodies following the same protocol and detected no labelling (Supp Fig. 7A). During the experiments, we identified the vessel type by the morphology, diameter and direction of flow, as done previously [44]. We observed VCAM-1 labelling surrounding the vessel wall in penetrating arterioles and capillaries but not in venules (Fig 7B and 7C). This labelling was “patchy”, i.e., not uniform throughout the vessel, as observed ex-vivo in other studies [65,66]. Both AD and WT mice expressed VCAM-1 in the vessel wall, which is consistent with previous studies of aged mice [65]. ICAM-1 was observed around the vessel wall in venules, capillaries, and penetrating arterioles (Fig. 7B and 7D).

**Figure 7.**
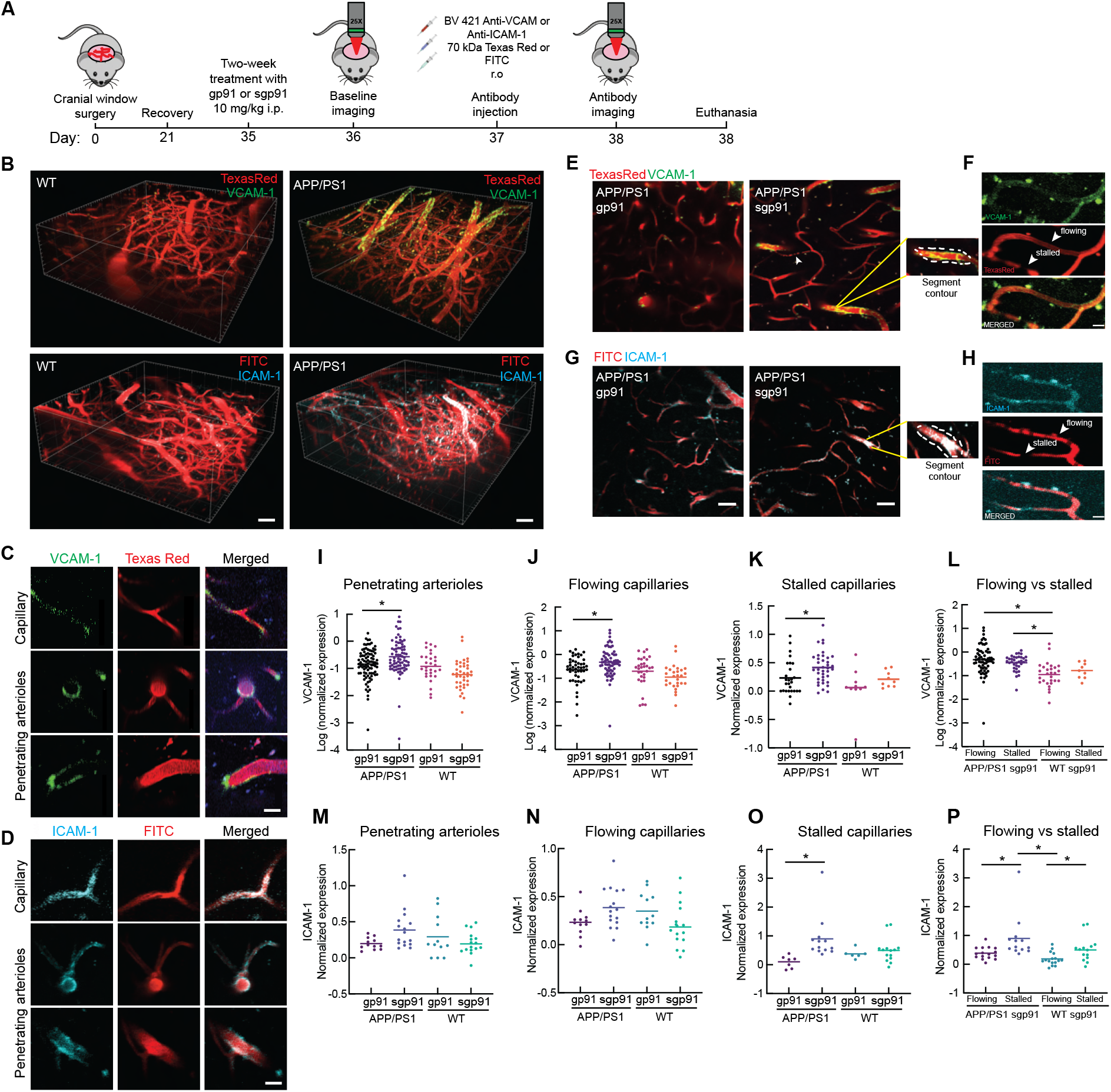
*In vivo* immunolabeling of vascular adhesion molecules in neocortical capillaries, penetrating arterioles, and venules in APP/PS1 mice. A) Experimental design, timeline and antibodies used. B) 3D renderings of vascular networks labeled with BV 421 anti-VCAM-1 and Texas Red, BV 421 anti-ICAM-1 and FITC. Differences between AD and WT are shown. Scale bar = 30 µm. C) Representative examples illustrating the labeling of penetrating arterioles and capillaries in APP/PS1 mice with BV 421 anti-VCAM-1 and Texas Red dextran for the vasculature. Scale bar = 20 µm. D) Representative examples illustrating the labeling of penetrating arterioles and capillaries in APP/PS1 mice with BV 421 anti-ICAM-1 and FITC dextran for the vasculature. Scale bar = 20 µm. E) 2D images of *in vivo* labeling of VCAM-1 and F) ICAM-1 in different types of vessels in AD mice treated with NOX2 inhibitor and control. Insets show zoomed images of representative penetrating arterioles that denotes the manually segmented contour of the vessel where the measurements were made. Scale bar = 50 µm. G) Example of a stalled vs flowing vessel labelled with VCAM-1 and Texas Red for the blood plasma. Scale bar = 10 µm. H) Example of a stalled vs flowing vessel labelled with ICAM-1 and FITC for the blood plasma. Scale bar = 10 µm. Quantification of VCAM-1 expression normalized by plasma labelling in I) Penetrating arterioles (one-way ANOVA, p < 0.05), J) Flowing capillaries (one-way ANOVA, p < 0.05), K) Stall-prone capillaries (Kruskal-Wallis test, p < 0.05) in WT and APP/PS1 mice treated with control or NOX2 inhibitor. L) Comparison of VCAM-1 expression from flowing vs stall-prone capillaries in APP/PS1 and WT controls (one-way ANOVA, p < 0.05). Data in I, J and L followed a skewed distribution; therefore, we used a logarithmic transform to normalize, visualize, and test for significance in the data. Quantification of ICAM-1 expression normalized by plasma labelling in M) Penetrating arterioles (one-way ANOVA, p < 0.05), N) Flowing capillaries (one-way ANOVA, p < 0.05), O) Stall-prone capillaries (Kruskal-Wallis test, p < 0.05) in WT and APP/PS1 mice treated with control or NOX2 inhibitor. P) Comparison of ICAM-1 expression from flowing vs stall-prone capillaries in APP/PS1 and WT controls (one-way ANOVA, p < 0.05).

We quantified the expression of these molecules in penetrating arterioles and capillaries by manually segmenting the vessel contour and measuring the integrated density of the antibody signal (Fig. 7E and 7F). We then normalized this signal to the plasma labelling, and pooled data by type of vessel, animal phenotype, and treatment. Stalled capillaries were also identified manually, as described elsewhere [29],.

Baseline imaging was performed before antibody administration to rule out potential effects of the antibody in modulating capillary stalls. We found no differences in capillary stalls pre and post antibody administration (Sup Fig. 7B), which indicates that fluorescently labelled antibodies do not interfere with neutrophil arrest.

VCAM-1 expression was significantly higher in penetrating arterioles, as well as flowing and stall-prone capillaries of AD mice, as compared to age-matched WT animals (Fig. 7I-L), and this upregulation was attenuated by a two-week treatment with NOX-2 inhibitor (gp-91). ICAM-1 showed a trend towards increased expression in penetrating arterioles and flowing capillaries and a significantly increased expression in stall-prone capillaries in AD mice, as compared to WT. This upregulation was also modulated by NOX2-inhibition (Fig. 7M-P). This data directly links vascular inflammation with amyloid-beta induced oxidative stress, specifically, in the penetrating arterioles, capillaries and obstructed vessels of the neocortex.

### 3.8 NOX2 is upregulated in the brain of AD patients as compared to age-matched controls

Staining of tissue from human brain banks (see Supplementary Table 1 for details on tissue sections) revealed upregulation of NOX2 in AD patients, as compared to controls (Supp Fig 8).

## 4. DISCUSSION

Increases in ROS-mediated cellular damage have been implicated in the pathogenesis of AD [67]. A dominant source of ROS in the brain are the various isoforms of NAPDH oxidase, with the NOX2 [68] and NOX4 [8] containing isoforms contributing most prominently to neurovascular dysfunction. Indeed, we observed an increased amount of NOX2 in both APP/PS1 mice, as compared to WT, and in AD patients, as compared to healthy controls, similar to previous findings [68]. In agreement with a previous study, we also observed that the administration of gp91-ds-tat reduced *Nox2* mRNA expression in the brain cortex of AD animals [69]. The mechanism by which this occurs is unknown, but might be a downstream result of the inactivation of the NADPH oxidase complex leading to reduced ROS-dependent activation of transcription factors that regulate pro-inflammatory gene expression, including *Nox2* [70].

Previous work has established that one consequence of NOX2-derived ROS in AD mouse models is the loss of normal neurovascular regulation. Cerebrovascular reactivity was significantly reduced in WT animals after topical superfusion of Aβ40, but improved with the administration of the NOX2 inhibitor gp91-ds-tat. Similarly, mice lacking the NOX2 subunit of NADPH oxidase were protected against Aβ-induced loss of cerebrovascular regulation [58]. In addition, NOX2 knock-out was shown to rescue behavioral deficits in the Tg2576 mouse model of APP overexpression [36]. Here, we show that these same NOX2-derived ROS also underlie the increased capillary stalling that causes reductions in resting state CBF in AD mouse models. Inhibiting NOX2 to reduce ROS, increasing peroxynitrate decomposition to reduce downstream DNA damage, or inhibiting PARP-1 to block the ROS damage response all decreased capillary stalling and increased CBF.

Consistent with previous findings [36], NOX2 inhibition led to improvements in performance on short-term memory tasks. Several factors could underlie this cognitive improvement. We have previously shown that an increase in CBF is followed by a rapid improvement in short term memory in APP/PS1 mice [71]. It is likely the CBF increase we observed after NOX2 inhibition similarly contributes to improved memory function. However, previous studies have shown that increased microglia activation and brain inflammation [72–74] correlate with poorer cognitive performance, and we found that microgliosis in the cortex and hippocampus decreased with NOX2 inhibition on the same timescale as the behavioral improvements, suggesting this decrease in brain inflammation may also contribute to the memory improvements. Consistent with previous studies of ROS inhibition in AD [58], we observed memory improvements in the absence of notable changes in parenchymal amyloid pathology. Taken together, these data suggest that other mechanisms independent of amyloid-beta, such as modulating blood flow and neuroinflammation, may be sufficient for improving memory.

To gain insight into the mechanisms that contribute to increased capillary stalling in AD mice, we performed unbiased RNA sequencing of cerebral microvessels to explore the underlying changes in gene expression that mediated these physiological effects. Knowing that NOX2 inhibition reduced capillary stalling, we looked for gene expression patterns that were changed in AD vs. WT and that were then normalized by NOX2 inhibition in the AD mice. We found enriched gene sets in AD microvessels related to inflammation, reactive oxygen species, and cell adhesion, when compared to WT controls, as seen in previous studies evaluating transcriptional changes in AD microvessels [64]. Furthermore, when analyzing enrichment in gene sets from AD gp91 vs AD sgp91, we found a downregulation in pathways related to inflammation and cell adhesion. Nonetheless, the lack of differentially expressed genes with NOX2 inhibition suggests that the effects on blood flow and memory function we found after treatment may be mediated mostly by physiological changes rather than transcriptional changes. For example, NOX2 inhibition reduces intracellular calcium in endothelial cells and decreases the production of superoxide anions and peroxynitrate [38], which may improve cellular function independent of gene expression changes.

Although subtle, we did find increased expression of cell adhesion pathways in AD that were downregulated by NOX2 inhibition. We further explored the relevance of these cell adhesion pathways and their role in capillary stalling with *in vivo* immunolabeling. Based on the study by Sipkins et al., [75], we used fluorescently conjugated antibodies against cellular adhesion molecules such as VCAM-1 and ICAM-1, with two-photon imaging to visualize the expression of these molecules in the mouse brain *in vivo*. We observed differential labelling of these adhesion molecules in capillaries, venules and penetrating arterioles, consistent with previous studies that have performed post-mortem labelling [65,66]. We found that VCAM-1 and ICAM-1 are mostly expressed in capillaries and penetrating arterioles, upregulated in AD animals, and downregulated with NOX2 inhibition. We saw no difference in VCAM-1 in stall-prone vs. flowing capillaries, suggesting that while VCAM-1 may be upregulated in AD, this is not unique in stall-prone vessels and so is not a sufficient condition to cause stalling, although it may still be necessary. ICAM-1, on the other hand, was more strongly upregulated in stall-prone capillaries in AD mice, as compared to flowing capillaries, suggesting it may play a direct role in neutrophil arrest in capillary segments. Other mechanisms of vascular dysfunction may be playing an important role in stalling together with increases in cellular adhesion molecules, such as pericyte constriction [8] and BBB leakage [30].

In summary, the expression of NOX2 is increased in APP/PS1 mice which, through downstream pathways that include peroxynitrate production and PARP-1 activation [38], leads to decreased CBF by increasing the incidence of capillary stalls, in addition to the loss of neurovascular regulation that has been previously reported [36,38,57,58]. Inhibition of NOX2 decreased capillary stalls, increased CBF, decreased brain inflammation, and rescued memory function. While direct ROS reduction has proven difficult or ineffective as an AD treatment in patients [76], future work to dissect the downstream mechanisms by which ROS reduction is leading to such improvements in mouse models could identify novel targets for therapy development.

## ACKNOWLEDGEMENTS

We would like to thank the brain banks who provided us with human brain tissue for this study: the NIH NeuroBioBank, the Harvard Brain Tissue Resource Center, Sepulveda Research Corporation, the Mount Sinai/JJ Peters VA Medical Center NIH Brain and Tissue Repository and the University of Maryland Brain and Tissue Bank. The confocal microscopy imaging data in this manuscript was acquired through the Cornell University Biotechnology Resource Center, with National Institutes of Health grant S10RR025502 funding for the shared Zeiss LSM 710 Confocal. Special thanks to Laura Berkowitz for help with statistical analyses. This work was supported by the American Heart Association Pre-Doctoral Fellowship grant 828650 (NERU), The Cornell Neurotech Mong Junior Fellowship (NERU), DFG German Research Foundation (OB), National Institutes of Health grants NS097805 (LP), NS114561 (TS), NS/HL37853 (CI), AG049952 (CBS), NS108472 (CBS), and BrightFocus Foundation grants A2017488S (CBS).

