## Supplementary Material for "Vascular oxidative stress causes neutrophil arrest in brain capillaries, leading to decreased cerebral blood flow and contributing to memory impairment in a mouse model of Alzheimer’s disease"

| **Area** | **Phenotype** | **Age** | **Sex** |
|---|---|---|---|
| Middle temporal (Brodmann area 21) | Unaffected Control | 69 | M |
| Middle temporal (Brodmann area 21) | Alzheimer's disease with late onset | 82 | F |
| Middle temporal (Brodmann area 21) | Alzheimer's disease, unspecified | 75 | F |
| Middle temporal (Brodmann area 21) | Unaffected Control | 79 | M |
| Middle temporal (Brodmann area 21) | Unaffected Control | 70 | F |
| Middle temporal (Brodmann area 21) | Alzheimer's disease, unspecified | 80 | F |
| Middle temporal (Brodmann area 21) | Unaffected Control | 84 | F |
| Middle temporal (Brodmann area 21) | Alzheimer's disease, unspecified | 62 | M |
| Middle temporal (Brodmann area 21) | Alzheimer's disease, unspecified | 89 | M |
| Hippocampus | Alzheimer's disease, unspecified | 82 | F |
| Hippocampus | Alzheimer's disease, unspecified | 74 | F |
| Hippocampus | Alzheimer's disease, unspecified | 76 | M |
| Hippocampus | Alzheimer's disease, unspecified | 84 | M |
| Hippocampus | Unaffected Control | 73 | F |
| Hippocampus | Personality change due to known physiological condition | 70 | M |
| Hippocampus | Unaffected Control | 62 | M |

**Supplementary Table 1.** Samples for human tissue staining.

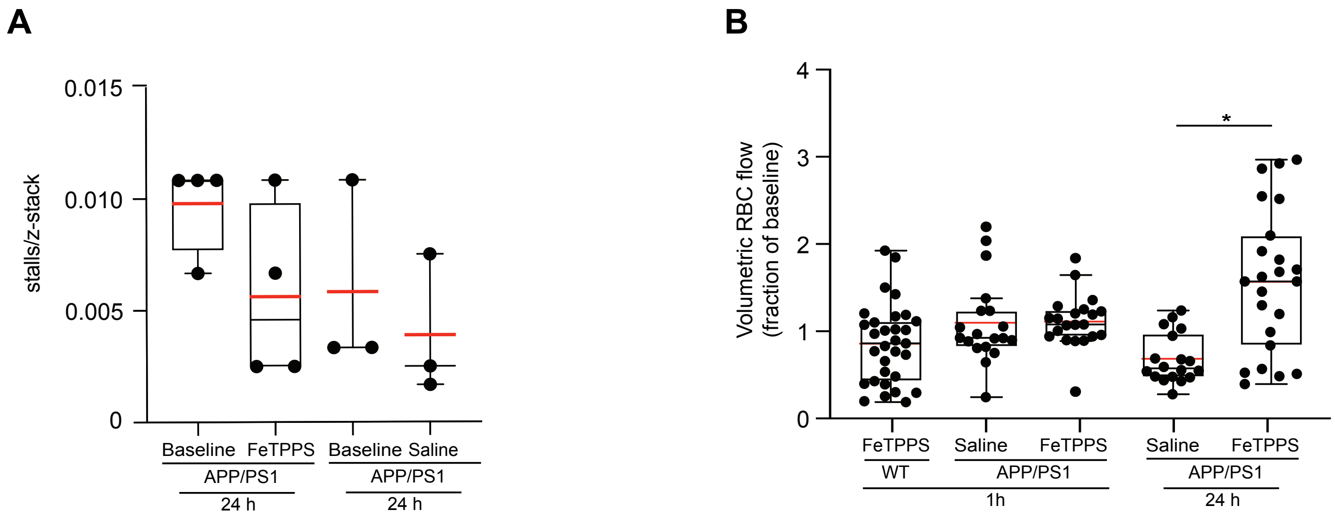


**Supplementary figure 1**. **Treatment with the peroxynitrate decomposition catalyst FeTPPS reduces capillary stalls and increases arteriole blood flow in APP/PS1 mice.** A) Fraction of capillaries with stalled blood flow in 10-11 months old APP/PS1 mice at baseline and after 24 h of treatment with FeTPPS (10 mg/kg, i.p.) (n = 4) or saline (n = 3). B) Penetrating arteriole blood flow volume, expressed as a fraction of baseline, from WT and APP/PS1 mice treated with the peroxynitrate decomposition catalyst FeTPPS (10 mg/kg, i.p.) or saline as control after 1h and 24h. Kruskal Wallis test, * $p < 0.05$.

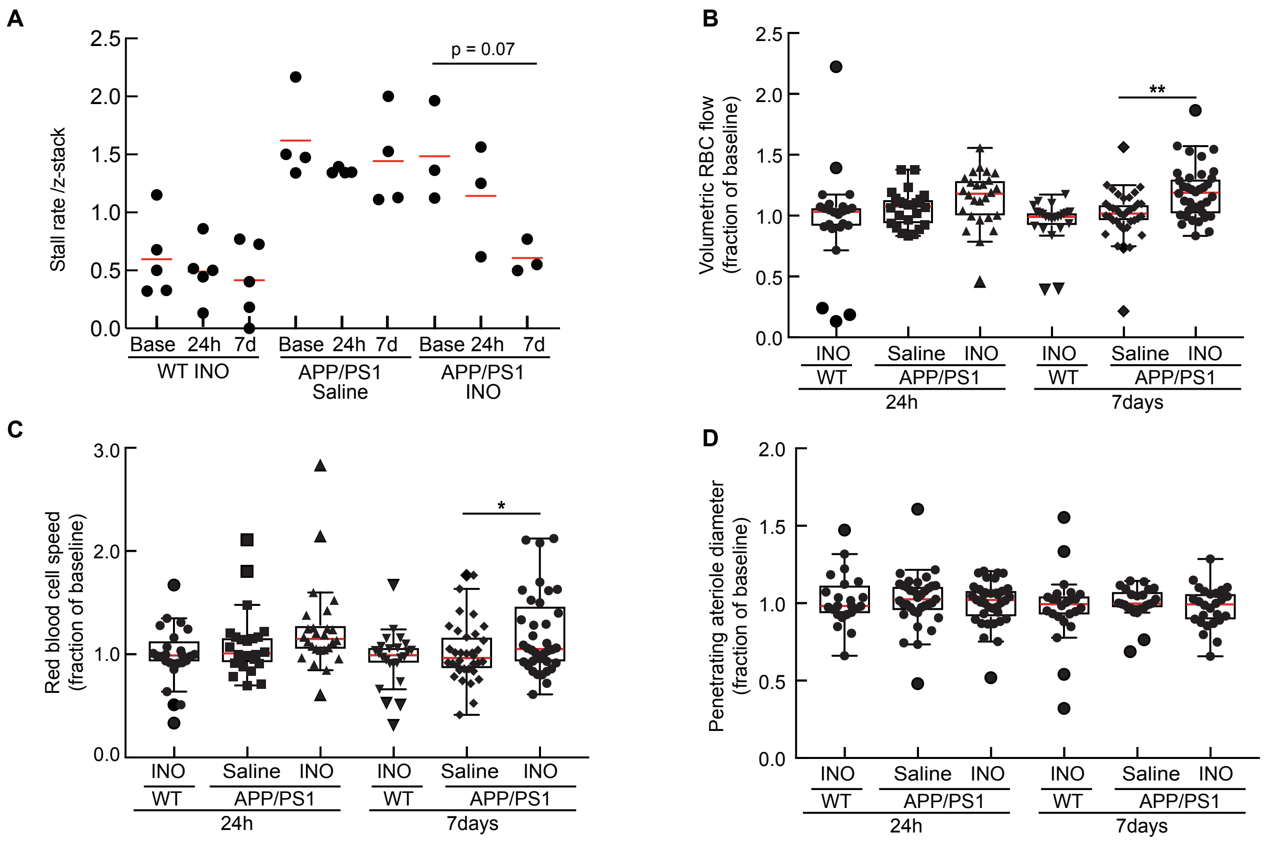


**Supplementary figure 2. Treatment with the PARP inhibitor 3-aminobenzamide reduces capillary stalls and increased penetrating arteriole flow in APP/PS1 mice.** A) Fraction of capillaries with stalled blood flow at baseline and after 24 hours and 7 days of treatment with the PARP inhibitor 3-aminobenzamide (10 mg/kg ;every other day for 7 days, i.p.) or saline in 10-11 months old APP/PS1 and WT mice. Penetrating arteriole B) blood flow volume, C) RBC speed, and D) vessel diameter expressed as a fraction of baseline, after 24 hours and 7 days of treatment with the PARP inhibitor 3-aminobenzamide or saline as a control in APP/PS1 and WT mice. Kruskal Wallis test with Tukey post-hoc comparisons, * $p < 0.05$; ** $p < 0.01$)

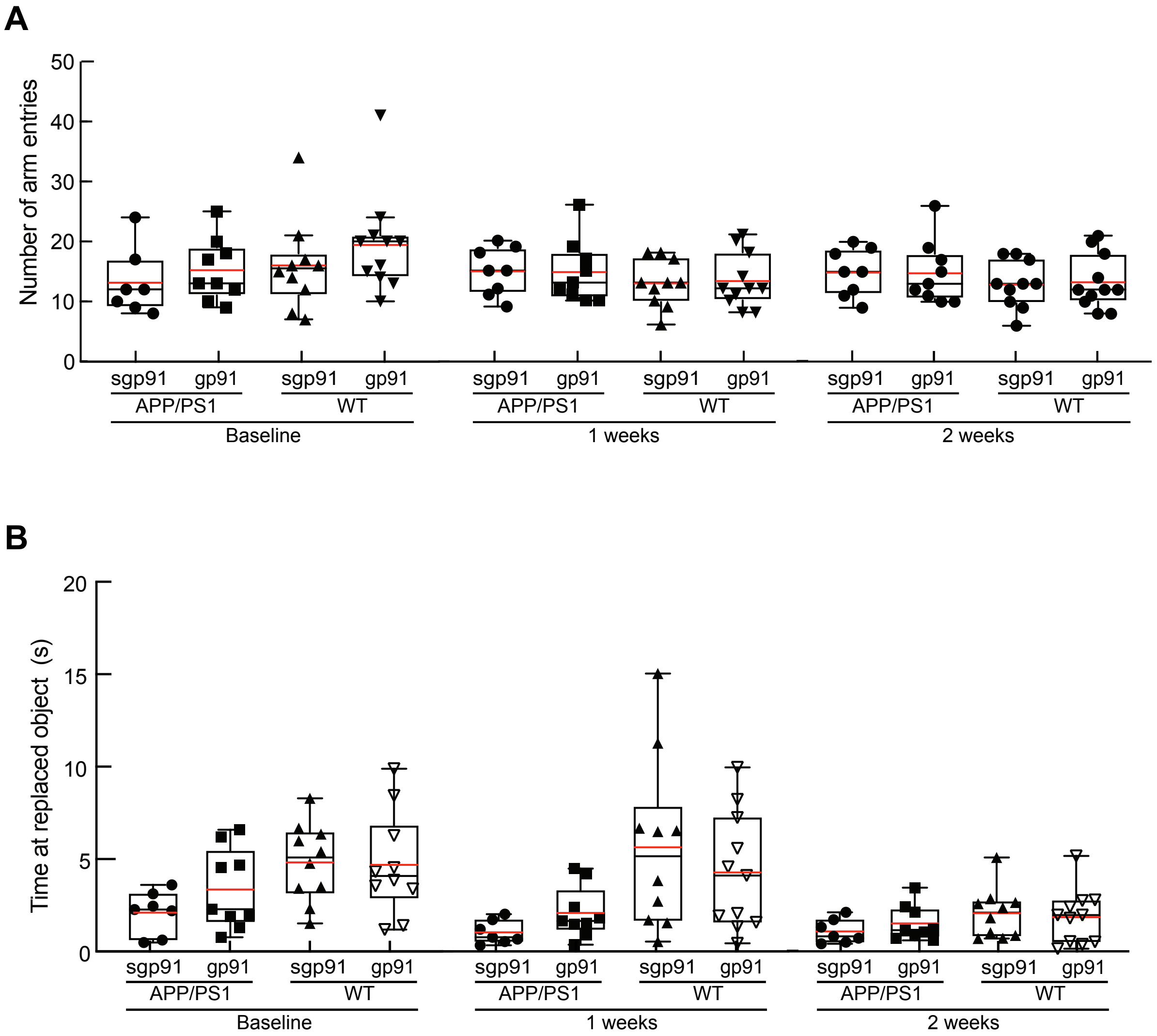


**Supplementary figure 3. Number of arm entries and time at moved object remain unchanged in behavioral tests across timepoints, suggesting continued engagement with the behavioral test.** A) Number of arm entries in the Y-maze and B) time at replaced object for the object displacement test at baseline, and after 1 and 2 weeks of treatment with the NOX2 inhibitor or scrambled control. No significant differences detected between groups with a one-way ANOVA test.

A

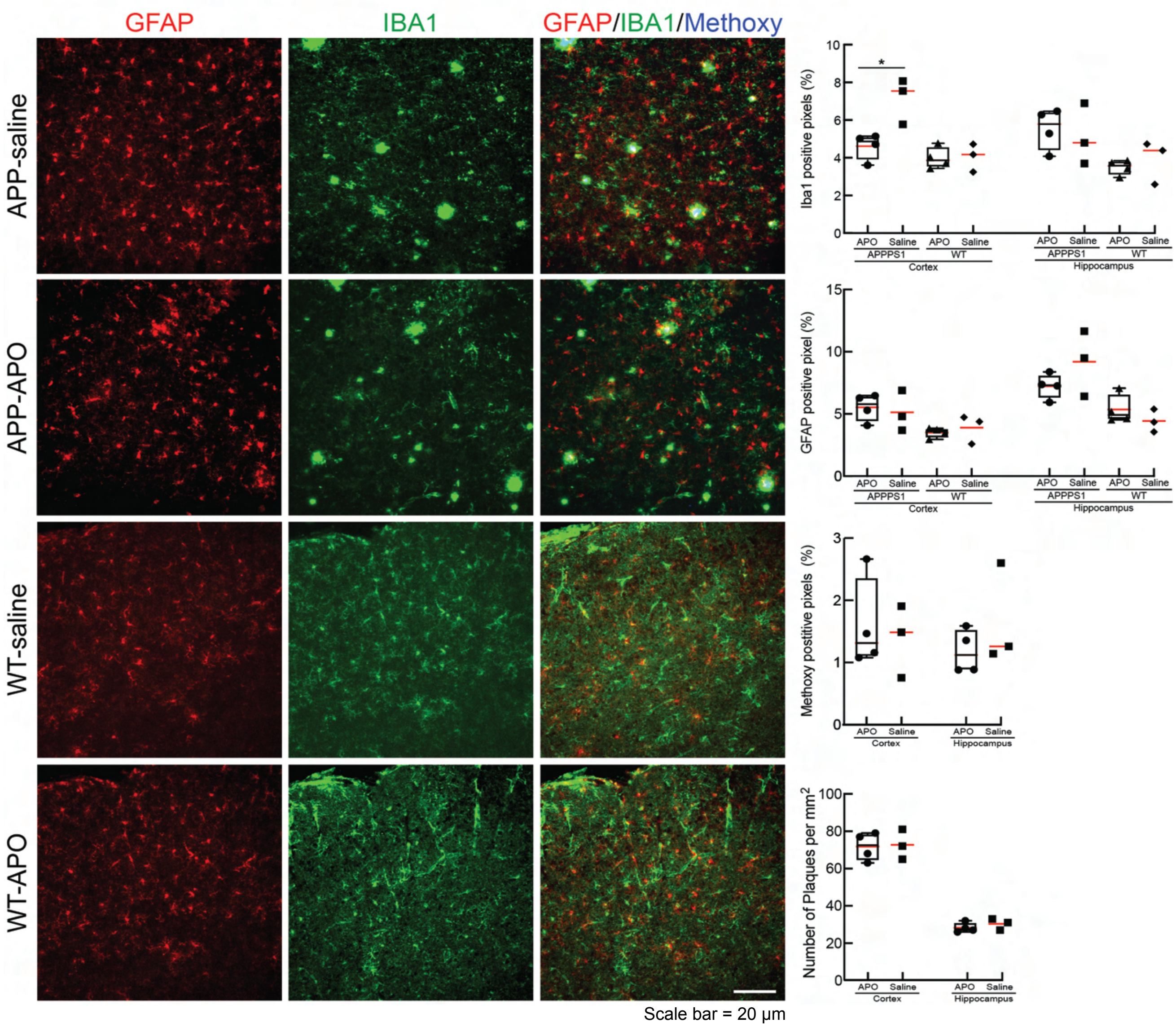


**Supplementary figure 4. Apocynin, a general NOX2 inhibitor, reduces the expression of IBA1 in the cortex of APP/PS1 mice.** GFAP, Methoxy X-04 and IBA1 expression in brain cortical tissue of APP/PS1 and WT mice (10-11 months old) treated with a 10 mg/kg dose of apocynin (via i.p.). A significant decrease in IBA1 positive pixels was detected after two-week treatment with apocynin. Kruskal Wallis test, *$p < 0.05$.

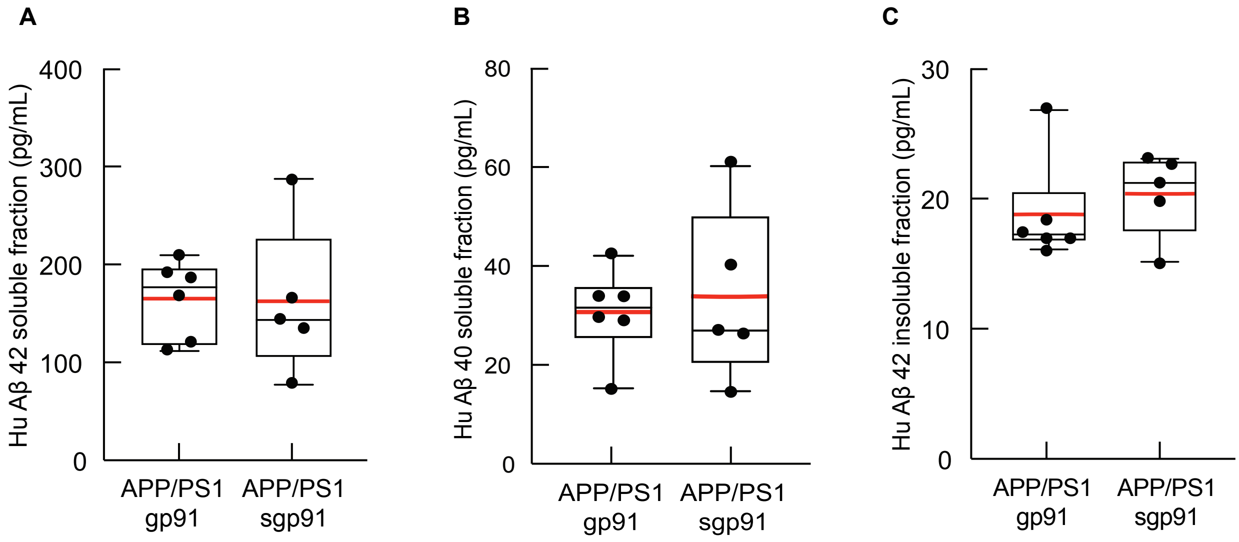


**Supplementary figure 5. Two weeks of NOX2 inhibition does not lead to changes in the concentration of amyloid beta species.** Results of Enzyme-linked immunosorbent assays (ELISA) of brain lysates from APP/PS1 mice treated with gp91-ds-tat or scrambled control sgp91-ds-tat. A) Soluble fraction of human Aβ 42, B) Soluble fraction of human Aβ 40 and C) Insoluble fraction of human Aβ 42. No significant differences were seen between groups.

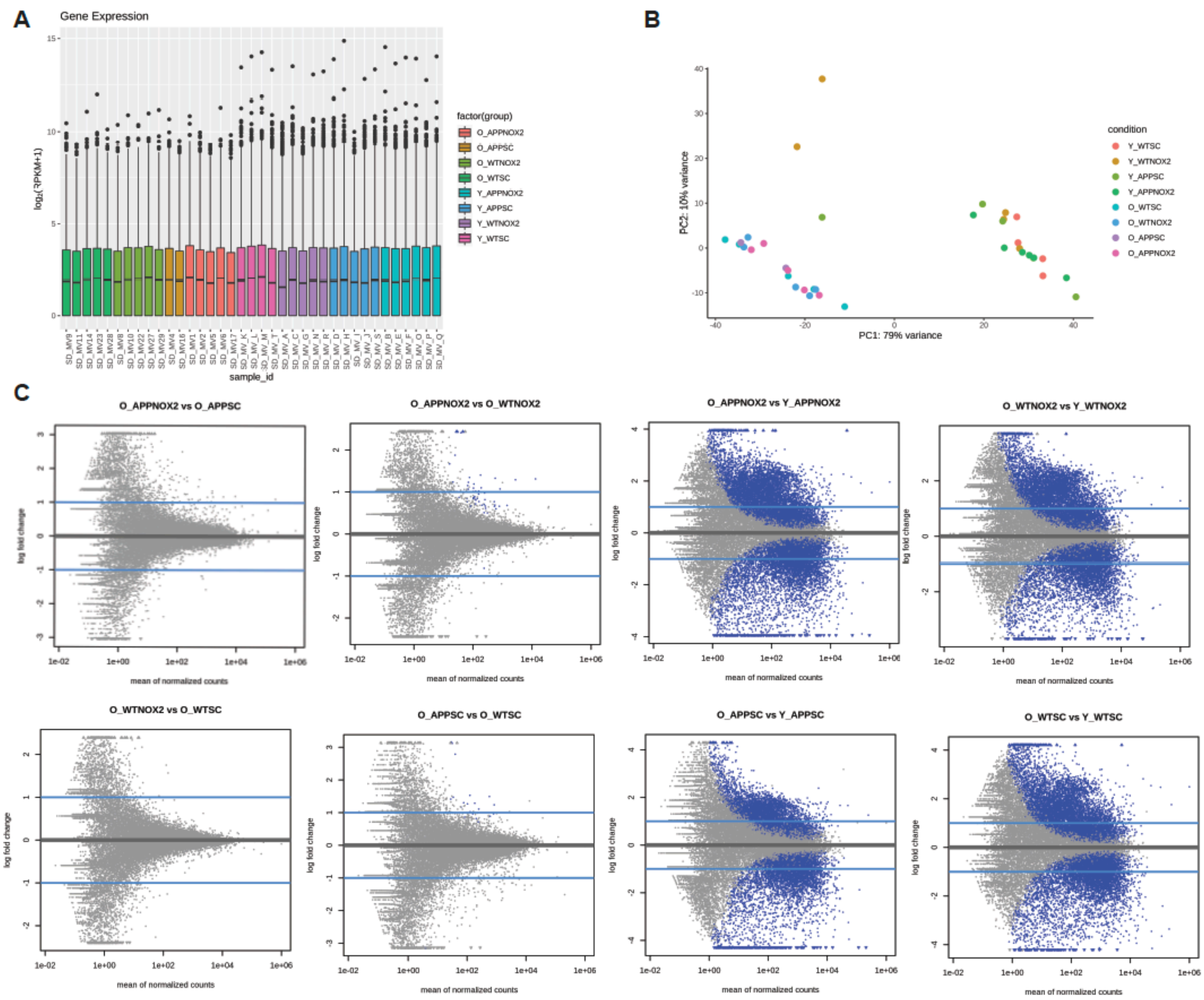


**Supplementary figure 6.** A) Number of reads (Log RPKM +1) per sample after RNA sequencing. B) PCA map of samples (O = old, Y = young, APP = APP/PS1, NOX2 = gp91, SC = sgp91). C) MA plots of differential gene expression between samples. Blue dots correspond to genes with padj < 0.1 (significant), log2FC threshold of 1. Significant differences were observed in old vs young groups, and APP/PS1 vs WT groups, but not between gp91 and sgp91.

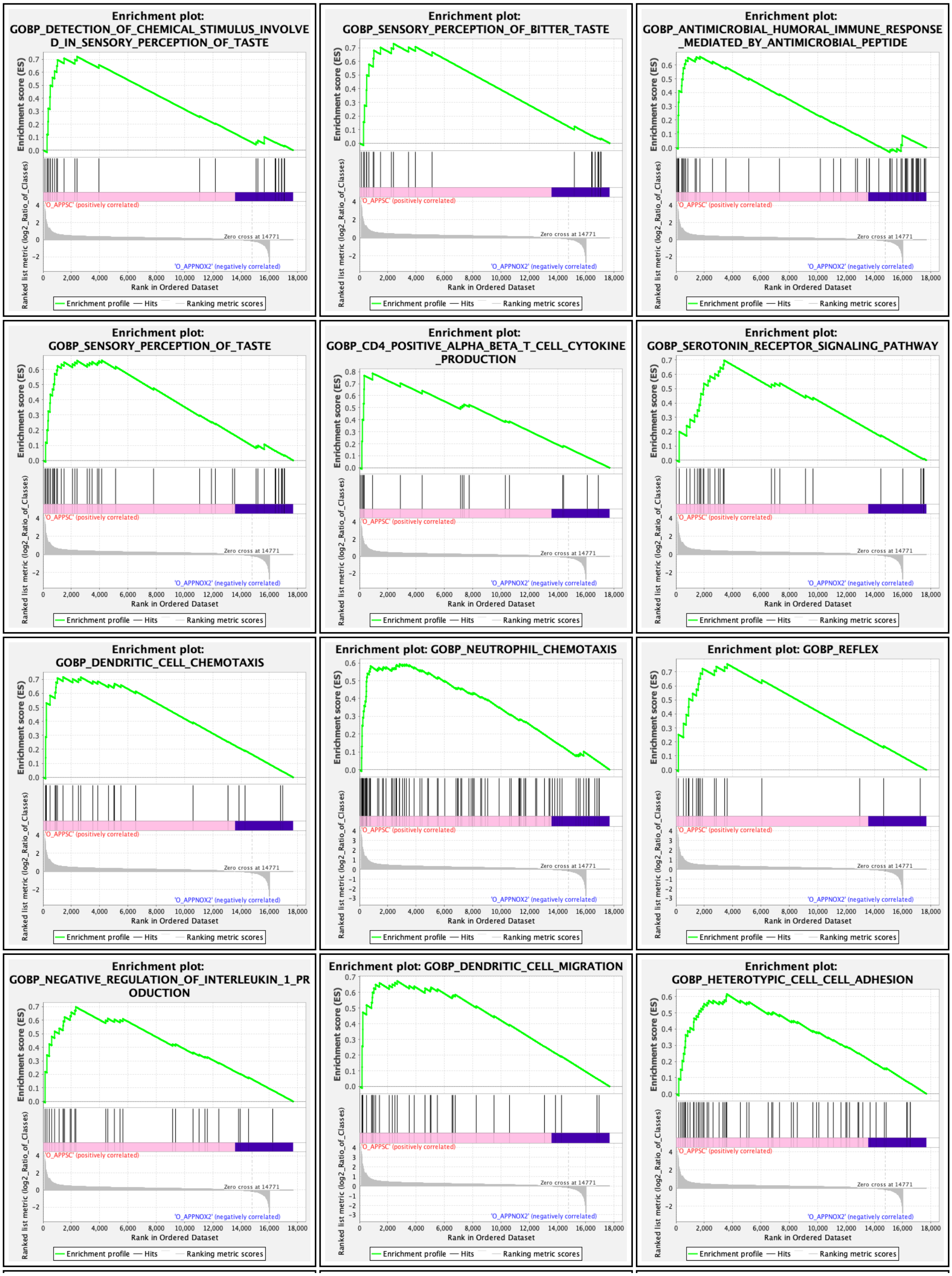

Enrichment plot: GOBP_DETECTION_OF_CHEMICAL_STIMULUS_INVOLVED_IN_SENSORY_PERCEPTION_OF_TASTE
Enrichment plot: GOBP_SENSORY_PERCEPTION_OF_BITTER_TASTE
Enrichment plot: GOBP_ANTIMICROBIAL_HUMORAL_IMMUNE_RESPONSE_MEDIATED_BY_ANTIMICROBIAL_PEPTIDE
Enrichment plot: GOBP_SENSORY_PERCEPTION_OF_TASTE
Enrichment plot: GOBP_CD4_POSITIVE_ALPHA_BETA_T_CELL_CYTOKINE_PRODUCTION
Enrichment plot: GOBP_SEROTONIN_RECEPTOR_SIGNALING_PATHWAY
Enrichment plot: GOBP_DENDRITIC_CELL_CHEMOTAXIS
Enrichment plot: GOBP_NEUTROPHIL_CHEMOTAXIS
Enrichment plot: GOBP_REFLEX
Enrichment plot: GOBP_NEGATIVE_REGULATION_OF_INTERLEUKIN_1_PRODUCTION
Enrichment plot: GOBP_DENDRITIC_CELL_MIGRATION
Enrichment plot: GOBP_HETEROTYPIC_CELL_CELL_ADHESION
Enrichment score (ES)
Ranked list metric (log2_Ratio_of_Classes)
'O_APPSC' (positively correlated)
Zero cross at 14771
'O_APPNOX2' (negatively correlated)
Rank in Ordered Dataset
Enrichment profile — Hits Ranking metric scores

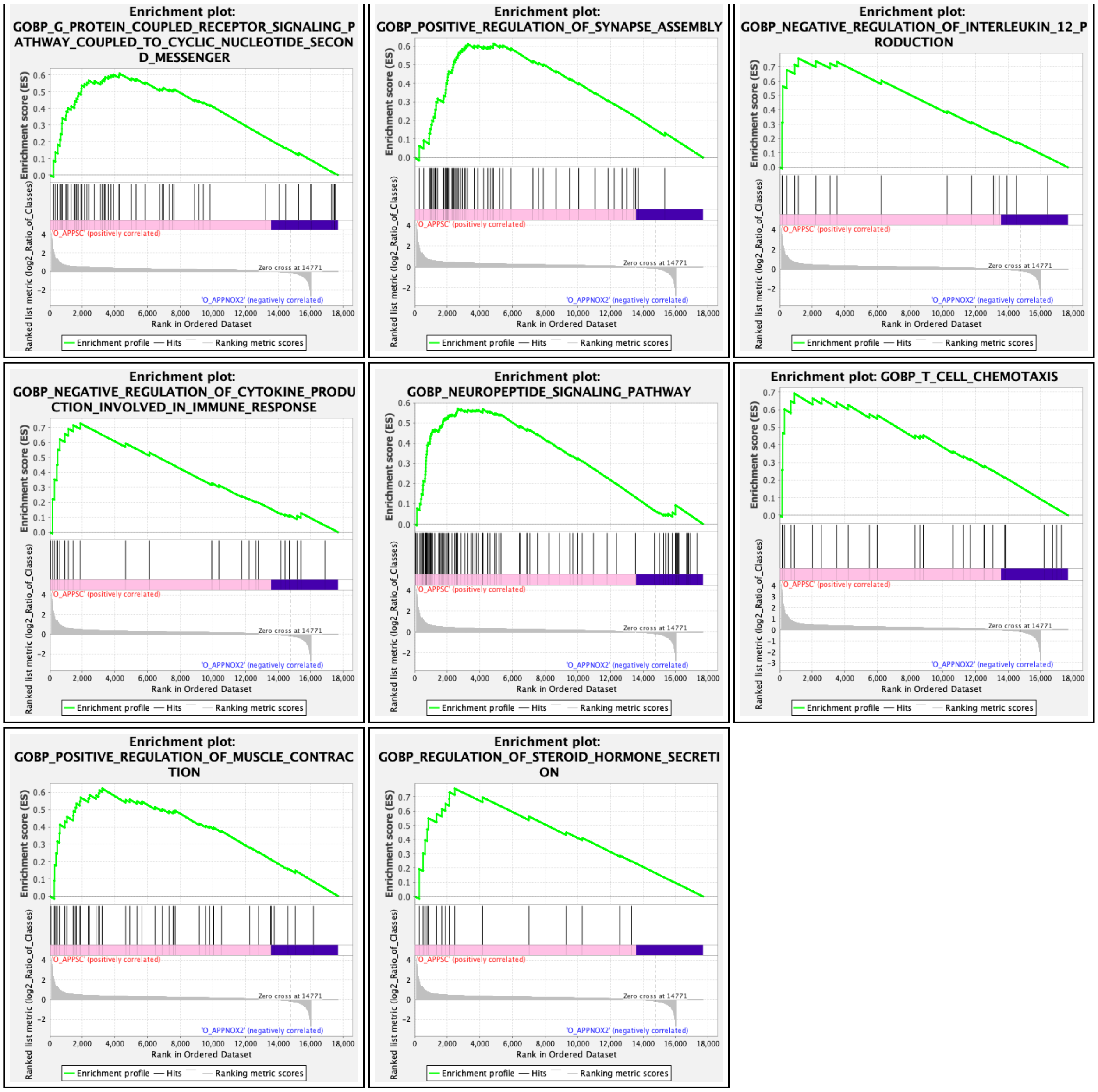


**Table 2.** Enrichment plots from GSEA analysis of AD sgp91 vs AD gp91 from the old mice group.

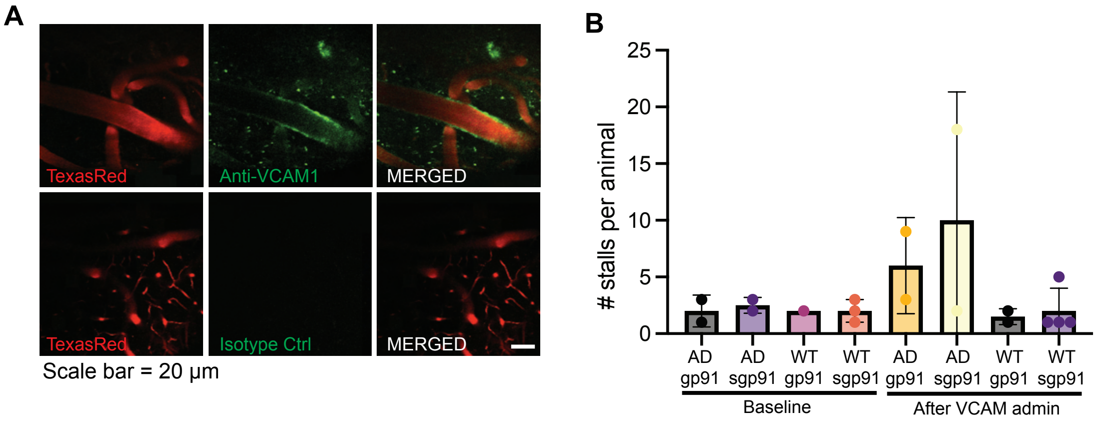


**Supplementary figure 7**. A) Representative two-photon images of cortical blood vessels labelled with anti-VCAM-1 , Isotype control and Texas Red for blood plasma. Positive labelling is observed with anti-VCAM1, but not with Isotype control. B) Average number of stalls detected per animal between groups. Baseline: AD gp91 (n=2), AD sgp91 (n=2), WT gp91 (n=1), WT sgp91 (n=3). After VCAM+P-selectin: AD gp91 (n=2), AD sgp91 (n=2), WT gp91 (n=2), WT sgp91 (n=3), Kruskal-Wallis test, $p < 0.05$. No significant differences were detected between groups.

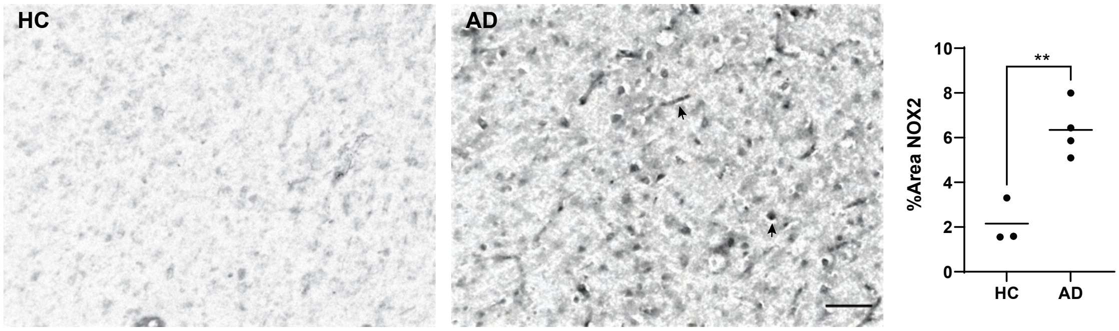


**Supplementary figure 8**. Expression of NOX2 (gp91-phox) in *post-mortem* cortical brain tissue of AD patients (n = 3) and healthy controls (n=4). Scale bar = 50 µm. Wilcoxon ranked t-test, $p < 0.05$.
